# A nutritional immunity blockade controls extracellular bacterial replication in *Legionella pneumophila* infections

**DOI:** 10.1101/2024.01.21.576562

**Authors:** Ascención Torres-Escobar, Ashley Wilkins, María D Juárez-Rodríguez, Magdalena Circu, Brian Latimer, Ana-Maria Dragoi, Stanimir S. Ivanov

**Affiliations:** Department of Microbiology and Immunology, Louisiana State University Health Sciences Center - Shreveport, Shreveport, LA 71130; Department of Molecular and Cellular Physiology, Louisiana State University Health Sciences Center - Shreveport, Shreveport, LA 71130; Innovative North Louisiana Experimental Therapeutics program (INLET), Feist-Weiller Cancer Center, Louisiana State University Health Sciences Center - Shreveport, Shreveport, LA 71130

**Keywords:** *Legionella pneumophila*, macrophage, monocyte, nutritional immunity, transferrin, extracellular replication, cell-mediated immunity, IFNγ

## Abstract

The accidental human pathogen *Legionella pneumophila* (Lp) is the etiological agent for a severe atypical pneumonia known as Legionnaires’ disease. In human infections and animal models of disease alveolar macrophages are the primary cellular niche that supports bacterial replication within a unique intracellular membrane-bound organelle. The Dot/Icm apparatus – a type IV secretion system that translocates ∼300 bacterial proteins within the cytosol of the infected cell – is a central virulence factor required for intracellular growth. Mutant strains lacking functional Dot/Icm apparatus are transported to and degraded within the lysosomes of infected macrophages. The early foundational work from Dr. Horwitz’s group unequivocally established that *Legionella* does not replicate extracellularly during infection – a phenomenon well supported by experimental evidence for four decades. Our data challenges the dogma in the field by demonstrating that macrophages and monocytes provide the necessary nutrients and support robust *Legionella* extracellular replication. We show that the previously reported lack of Lp extracellular replication is not a bacteria intrinsic feature but rather a result of robust restriction by serum-derived nutritional immunity factors. Specifically, the host iron-sequestering protein Transferrin was identified as a critical suppressor of Lp extracellular replication in an iron-dependent manner. In iron-overload conditions or in the absence of Transferrin, Lp bypasses growth restriction by IFNγ-primed macrophages though extracellular replication. It is well established that certain risk factors associated with development of Legionnaires’ disease, such as smoking, produce a chronic pulmonary environment of iron-overload. Our work indicates that iron-overload could be an important determinant of severe infection by allowing Lp to overcome nutritional immunity and replicate extracellularly, which in turn would circumvent intracellular cell intrinsic host defenses. Thus, we provide evidence for nutritional immunity as a key underappreciated host defense mechanism in *Legionella* pathogenesis.

## Introduction

*Legionella* species are environmental Gram-negative γ-proteobacteria that parasitize and replicate within free-living unicellular eukaryotes.^1,2^ *Legionella pneumophila* (Lp) is the leading cause of a lower respiratory infection known as legionellosis that produces an atypical pneumonia with a significant disease burden.^3^ Inhalation and deposition of aerosolized bacteria in the alveoli triggers the early flu-like symptoms of legionellosis - including dry cough, chills and fever – but disease progression varies from a mild self-limiting syndrome (aka Pontiac fever) to a debilitating severe pneumonia (aka Legionnaires’ disease), which can be lethal without appropriate antibiotics treatment.^4,5^ Outbreaks from a single clinical isolate frequently manifest in both the mild and the severe disease forms^6–13^, implicating host-determinants as important drivers of disease progression. History of smoking, old age and ongoing immunosuppression therapy represent key risk factors for progression of legionellosis towards Legionnaires’ disease^4,5^, however the mechanistic links remain unresolved.

In human infections and animal models of disease, alveolar macrophages are the primary cellular niche for bacterial replication.^14–18^ When internalized by phagocytes Lp rapidly remodels the early phagosome into a unique endoplasmic reticulum (ER)-like organelle, which within 4 hours post uptake matures to a vacuolar niche that supports robust bacterial replication.^19–22^ The critical virulence factor for intracellular replication conserved in all Legionellaceae is a type IVb secretion system (T4SS), referred to as the Dot/Icm apparatus, which translocates ∼300 bacterial proteins into the cytosol of the infected cell.^23,24^ Mutant strains lacking functional T4SS are transported to and degraded within lysosomes of infected macrophages.^25^ Although axenic growth conditions have been developed, *Legionella* does not replicate extracellularly during infection;^26,27^ thus, mutants that cannot grow intracellularly (such as T4SS− strains) are avirulent in both animal and tissue culture models of infection.^28–31^ Convincing experimental data from numerous labs over the years demonstrate that T4SS– bacteria are gradually eliminated when cultured with either human or murine macrophages or unicellular amoebae, whereas the wild-type strain grows several orders of magnitude – consistent with the idea of ‘obligate’ intracellular lifestyle during infection.^27,29,31–34^ Moreover, *Legionella* is fastidious due to auxotrophy for several proteogenic amino acids, including cysteine, which are acquired intracellularly from the host cell and partially explain the inability of the bacteria to replicate extracellularly in cellular infection models.^35–39^ Since the foundational work by Dr. Horowitz 40 years ago, the doctrine in the field is that *Legionella* under infection conditions replicates strictly intracellularly.^40^ Indeed, *Legionella* has biphasic developmental program typical of obligate vacuolar bacterial pathogens^41–46^ where a microorganism cycles between two phenotypically distinct forms each with a unique gene expression profile.^47–51^ A motile highly infectious ‘transmissive’ form, which upregulates the T4SS, is the predominant extracellular form detected during infection and in late exponential/early stationary phase in axenic growth cultures.^51^ The transition into a non-motile poorly infectious ‘replicative’ form, which is indistinguishable from the bacterial population during exponential growth in axenic cultures, occurs intracellularly within the vacuolar niche.^51^ Thus, in terms of its lifecycle during infection *Legionella* resembles obligate vacuolar bacterial pathogens.

An inflammatory response coordinated by multiple innate immune cell types results in IFNγ production, which resolves legionellosis by reprograming macrophages into a phenotypic state that restrict intracellular bacterial replication.^6,40,52–56^ IFNγ-stimulated primary human alveolar and monocyte-derived macrophages limit Lp intracellular replication *in vitro*^52,53^ through a variety of cell autonomous defense mechanisms in part impinging on LCV rupture and host cell death.^57–62^ IFNγ secretion is required for legionellosis resolution in animal models, emphasizing the importance of IFNγ effector functions in Lp pathogenesis.^55,56^ Yet, leukocytes from recovered Legionnaires’ disease patients secreted IFNγ when stimulated with Lp similarly to leukocytes isolated from the control cohort indicating that differences in IFNγ responses alone cannot account for disease progression.^63^ Thus, the determinants that allow a mild self-resolving legionellosis (aka Pontiac fever) that is well-controlled by innate immunity and IFNγ to progress into a severe atypical pneumonia are unclear.

Host defense mechanisms that sequester various trace metals and other micronutrients from invading pathogens, thus limiting pathogen replication and disease severity, are collectively known as nutritional immunity.^64–66^ Iron is essential for life and therefore is under strict nutritional immunity protection where high affinity iron-binding carriers, such as Transferrin and Lactoferrin, limit bioavailable iron in the extracellular milieu.^67^ Imbalance in systemic iron homeostasis - a complex tightly regulated process - increases susceptibility to bacterial infections especially under systemic iron-overload brought by saturation of host iron-carrier as a result of hereditary genetic disorders or high dietary iron consumption.^68–72^ Macrophages control bioavailable iron locally and systemically by regulated uptake of iron-carrier proteins via cell-surface receptors and secretion through Ferroportin (FPN1) – the only known iron exporter protein.^73,74^ During inflammation FPN1 is downregulated, which favors intracellular iron accumulation and a decrease in extracellular iron bioavailability.^69,75^ Nutritional immunity in legionellosis is poorly understood; nevertheless, under iron overload conditions IFNγ-primed macrophages fail to restrict Lp growth *ex vivo* suggesting that compromised nutritional immunity impacts cell autonomous responses to Lp.^76,77^ The molecular mechanism driving this phenotype remains unresolved. Here, we provide important novel insight in the cooperation between nutritional and cell mediated immunity for the restriction of Lp growth.

## Results

### Legionella mutant strains lacking a functional Dot/Icm secretion apparatus replicate in the presence of human macrophages under serum-free conditions

We measured the intracellular growth kinetics of different Lp strains in human macrophage infections to investigate the connection between serum derived nutrients and bacterial replication. To avoid a pyroptotic cell death response triggered by cytosolic delivery of flagellin,^78^ which can convolute data interpretation in infection-based growth assays, we used a flagellin clean deletion mutant. We engineered two distinct *L. pneumophila* Philadelphia-1 derived strains – Lp01 and JR32 – to bioluminescence by inserting the LuxR operon (*luxC luxD luxA luxB luxE*) from *Photorhabdus luminescens* on the bacterial chromosome under the constitutive *IcmR* promoter using two different genetic approaches^79,80^ (SFig. 1a). For both strains the bioluminescence output increased exponentially during logarithmic growth in axenic cultures and peaked as the bacteria entered stationary phase (SFig. 1b-c).^79,80^ The Lp growth kinetics with human U937 macrophages peaked at 48hrs for both JR32 and Lp01 and the curves were comparable regardless of serum availability (Fig. 1 a-b). As negative controls in these assays, we used *dotA* clean deletion strains on the respective backgrounds, which lack functional Dot/Icm apparatus and have been extensively used in the *Legionella* field as negative growth controls because *dotA* mutants cannot replicate intracellularly.^25,32^ In the presence of serum, the bioluminescence signal in macrophage infections with the *dotA* mutants decreased sharply and remained below the level produced by the inoculum, consistent with the demonstrated inability of *dot*/*icm* deletion strains to replicate intracellularly (Fig. 1a-b). Unexpectedly, in infections under serum-free (SF) conditions *dotA* strains appear to replicate similarly in magnitude to the parental Dot/Icm+ strains, albeit with delayed kinetics (Fig. 1a-b). Area under the curve analysis from at least 9 biological replicates demonstrates the growth restriction-capacity of serum specifically towards the *dotA* mutant. The unexpected growth of the *dotA* mutant in SF infections was confirmed by an alternative Colony Forming Units (CFUs) growth assay, in which bacteria were collected at distinct times post infection and subsequently were serially diluted, plated and enumerated (Fig. 1d). When serum was present, Lp01 Δ*dotA* CFUs decreased 100-fold over the course of the infection, whereas in the absence of serum 100-fold increase in CFUs occurred for the same time period (Fig. 1d). Unlike Dot/Icm-strains, Dot/Icm+ strains fail to grow on CYE plates supplemented with 150mM NaCl due to ion imbalance.^81^ Therefore, to confirm that *dotA* bacteria did not regain Dot/Icm functionality in SF infections, eight recovered colonies from every infection condition at each time point were randomly selected and replica plated under high- and low-salt conditions. The results clearly demonstrate that CFUs recovered from Lp01 Δ*dotA* infections remained salt resistant regardless of serum supplementation, confirming the Dot/Icm-genotype (Fig. 1d).

**Fig 1.**
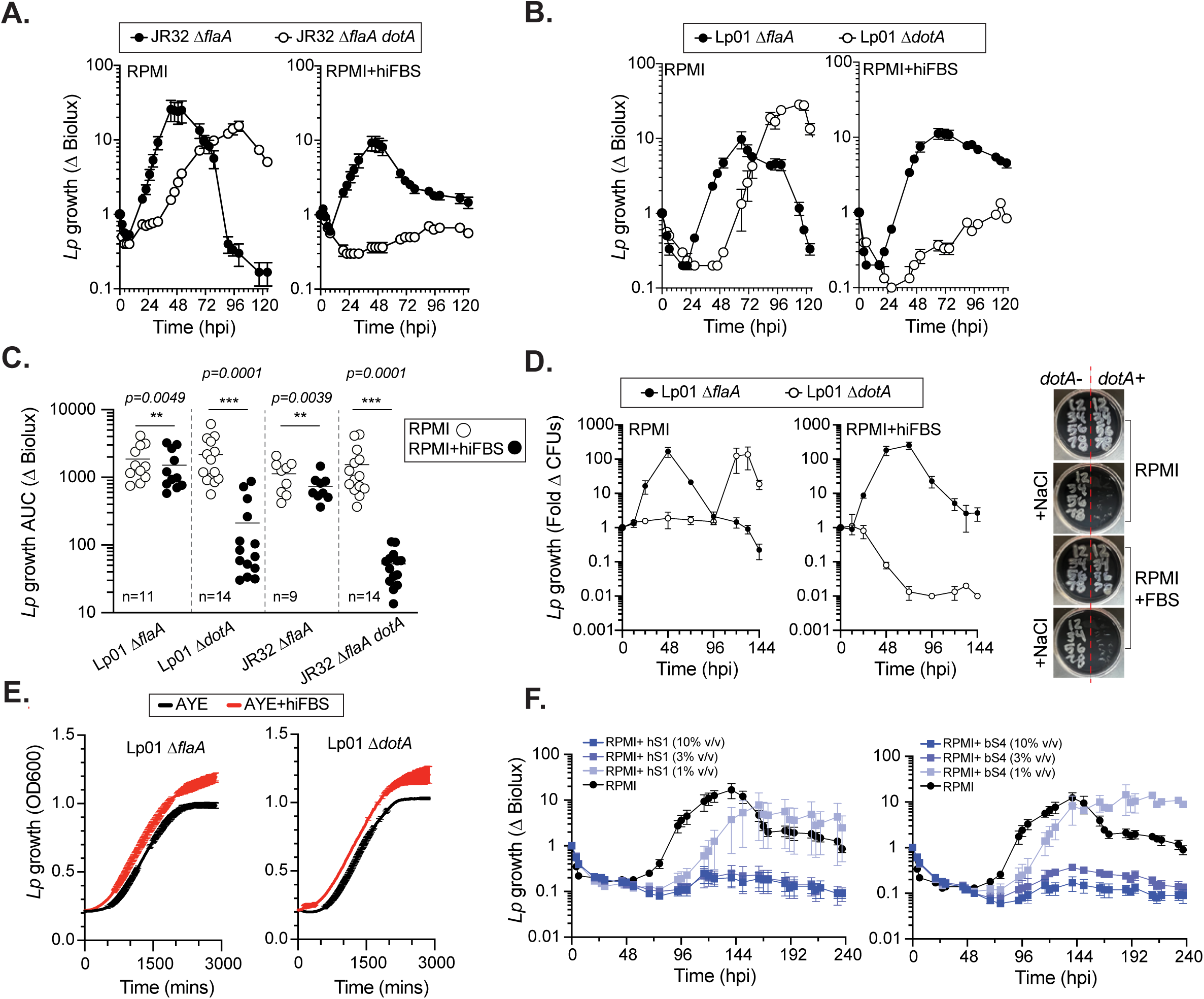
Effects of serum supplementation on Lp growth in human macrophage infections. **(A-D)** Growth kinetics of DotA+ and DotA– Lp in U937 MFs cell culture infections. Bacterial growth was measured at discrete time-points via bioluminescence output **(A-C)** or recovered CFUs **(D)** for mutant strains on the JR32 **(A, C)** and the Lp01 **(B-D)** backgrounds as indicated. **(C)** Area under the curve (AUC) analysis of Lp growth kinetics in the presence vs. absence of hiFBS for each infection condition. Each data point represents the AUC calculated from the growth curve in a single biological replicate as shown in A and B. Statistical significance was determined by non-parametric Wilcoxon matched-pairs T-test and the respective p-values are indicated; n = number of biological replicates. **(D)** In the CFU growth assays for every timepoint eight of the recovered colonies were selected at random and replica plated on CYA agar with or without 150mM NaCl. **(E)** Axenic growth curves of the indicated strains in AYE liquid cultures in the presence/absence of 10% (v/v) hiFBS were generated from OD_600_ measurements every 10min over the indicated time period. Each time point represents an average of a technical triplicate. **(F)** Growth kinetics of JR32 Δ*flaA dotA* in U937 MFs cell culture infections supplemented or not with different amounts of heat-inactivated human (hS1) or heat-inactivated bovine serum (bS4). **(A-B, D-F)** Averages from three technical replicates ± StDev are shown. At least three biological replicates were completed for each experiment.

Because the growth restriction phenotype was specific for the *dotA* mutants, we investigated whether *dotA* deletion sensitized Lp to a yet unidentified bacteriocidal component of the hiFBS used in the previous infection assays, which would explain the selective growth-restriction phenotype. However, the *dotA* mutant growth rate in axenic cultures was not reduced upon addition of hiFBS (Fig. 1e), which is inconsistent with the presence of an anti-microbial activity in the serum. Sera from human and bovine origin blocked *dotA* replication in cellular infections in a dose-dependent manner (Fig. 1f). A panel of 12 commercially-sourced sera used in macrophage infections (SFig. 2a) and axenic growth assays (SFig. 2b) revealed that while sera blocked *dotA* replication with various potency in cellular infections, Lp growth in axenic cultures was not affected. Thus, we conclude that serum-mediated growth restriction by a yet unidentified mechanism is required for the previously reported inability of the *dotA* mutant to growth in macrophage infections.^25,32^

The *dotA* growth in macrophage infections was puzzling because this strain, similar to other *dot*/*icm* deletion mutants, is readily cleared via lysosomal degradation after internalization by the host cell. Thus, we investigated whether the growth phenotype is common to other mutant strains lacking a functional Dot/Icm system. To this end, we engineered single gene knockout strains lacking either *dotB* or *dotH* - two genes essential for a functional Dot/Icm apparatus. DotH participates in the formation of the outer membrane cap and the periplasmic ring of the Dot/Icm T4SS core complex,^82,83^ whereas the DotB is the energy-generating ATPase that facilitates translocation of effector proteins.^84,85^ The growth kinetics of *dotB* and *dotH* mutant strains in macrophage infections phenocopied the data from the *dotA* deletion strains (Fig. 2a) and all *dot*/*icm* deficient strains grew under high salt concentrations (Fig. 2b). Deletion of *dotB* in *Legionella jordanis* (SFig. 1a) - another *Legionella* species that encodes a Dot/Icm system and replicates intracellularly^86^ - produced a strain capable of replication in macrophage infections under SF conditions (SFig. 3a). Similar to the Lp *dotA* deletion strains, the Lj *dotB* mutant grew in axenic cultures regardless of serum supplementation (SFig. 3b). Thus, we conclude that serum from bovine and human origins restrict the replication of *dot*/*icm* mutants from *L. pneumophila* and *L. jordanis* specifically in macrophage infections but not in axenic growth cultures.

**Fig 2.**
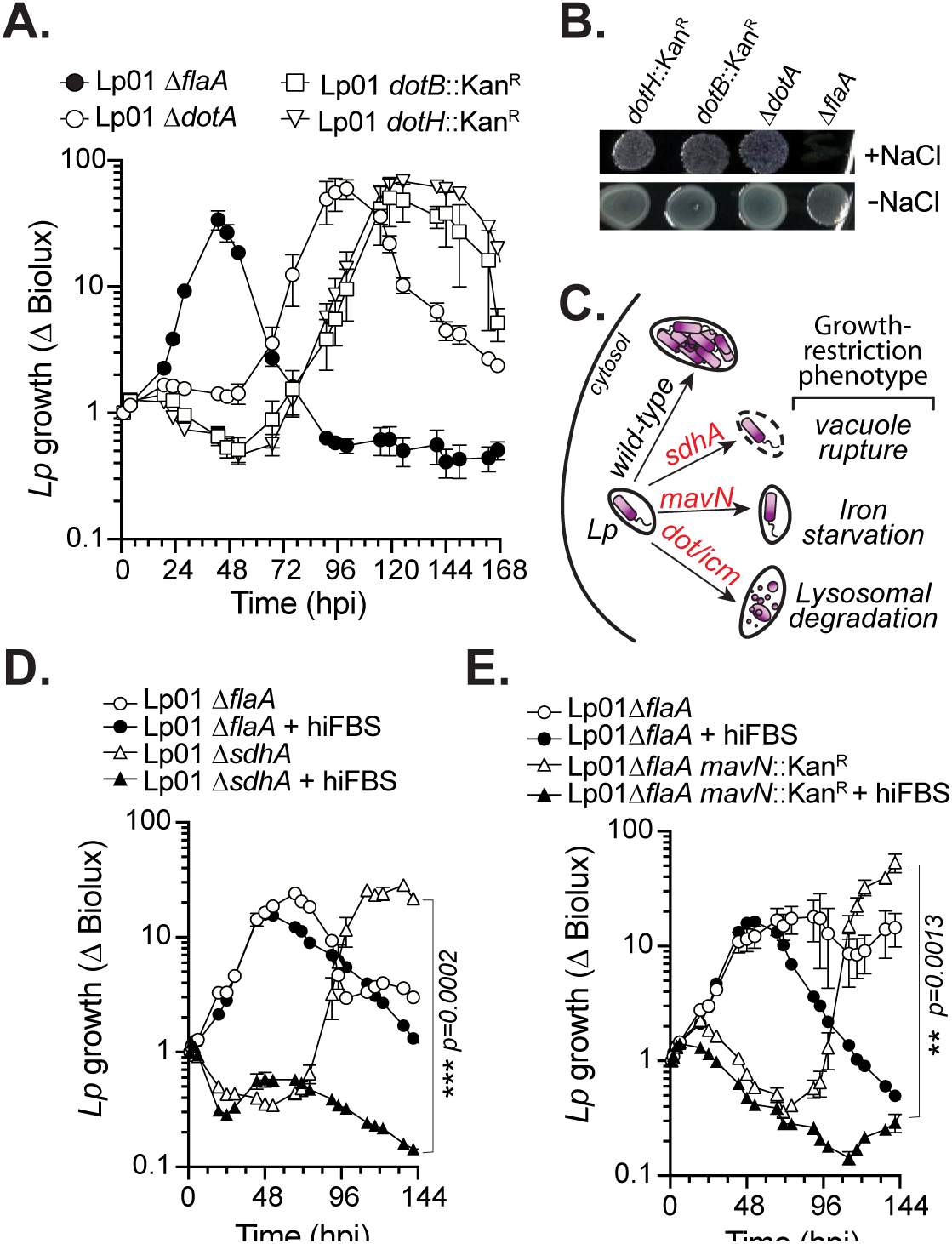
Serum withdrawal restores the growth of several Lp01 mutants with distinct intracellular growth defects. **(A)** Growth kinetics of different Lp01 Dot/Icm mutant strains with U937 macrophages under SF conditions. **(B)** Lp01 Dot/Icm mutants grow in the presence of 150mM NaCl. **(C)** Lp mutant strains (T4SS−, *mavN* and *sdhA*) exhibit intracellular growth-restriction phenotype due to distinct mechanisms – lysosomal degradation, iron starvation and vacuolar rupture. **(D-E)** Growth kinetics of the Δ*sdhA* **(D)** and *mavN*::Kan^R^ **(E)** mutants with U937 macrophages in the presence or absence of serum. Averages from three technical replicates ± StDev are shown. Statistical significance was determined by one-way ANOVA and the respective p-values are indicated. **(A-E)** At least three biological replicates were completed for each experiment.

### Serum restricts L. pneumophila extracellular replication in macrophage and monocyte infections

In light of the well-documented inability of (i) *dot*/*icm* mutants to avoid lysosomal degradation^25,32^ and (ii) Lp to replicate extracellularly during infection,^40^ we sought to determine which of the two postulates is affected by serum availability during infection. We took advantage of two other Lp knockout strains – *sdhA* and *mavN* – which fail to replicate specifically in macrophage infections because of distinct intracellular growth defects (Fig. 2c) but otherwise grow normally in axenic cultures.^87,88^ SdhA is a Dot/Icm effector protein involved in the early stages of LCV remodeling^89^ and *sdhA* deletion results in rapid destabilization of the Lp-occupied phagosome, which triggers a pyroptotic host-cell death response trapping the bacteria within the macrophage corpse thus restricting bacterial replication.^87^ Conversely, MavN is multi-transmembrane domain iron transporter translocated by the Dot/Icm apparatus that delivers ferrous iron across the LCV membrane.^88,90,91^ The LCVs occupied by *mavN* deletion mutants are remodeled and traffic normally, however bacterial replication is arrested due to iron starvation.^88^ Indeed, mutants lacking *sdhA* (Fig. 2d) or *mavN* (Fig. 2e) did not replicate in macrophage infections when serum was present; however, both strains grew with kinetics similar to the *dot*/*icm* deletion strains under SF conditions (Fig. 2a). Taken together, these data from infections with Lp mutants incapable of intracellular growth points to extracellular replication as a putative mechanism facilitating bacterial growth under SF conditions. This notion challenges the prevailing dogma that Lp cannot grow outside of the host cell during infection.^40^

Next, we set out to test directly whether Lp can replicate extracellularly during infection. To this end, we focused on monocyte infections because Lp entry in monocytes requires opsonization and FcR-mediated phagocytosis, which allows us to dictate bacteria location during infection. Using microscopy, we confirmed that U937 macrophages phagocytosed both opsonized and non-opsonized Lp efficiently, whereas U937 monocytes strictly internalized opsonized bacteria (Fig. 3a). Importantly, serum supplementation did not alter the Lp internalization determinants for either cell line (Fig. 3a). In monocyte infections, opsonized Lp initiated bacterial replication quickly and achieved maximal growth at ∼36 hpi (Fig. 3b) with kinetics similar to intracellular growth in macrophages (Fig 2a). Non-opsonized Lp also replicated in monocytes infections but with an obvious delay where growth was initiated at ∼ 48hpi and peaked at ∼ 96hpi (Fig. 3b). The growth kinetics of non-opsonized Lp in monocyte infections (Fig. 3b) and *dot/icm* mutants in both macrophage and monocyte infections (Figs. 2a and 3b) were comparable, which would be predicted if growth determinants in all circumstances were similar. Importantly, non-opsonized Lp replicated in monocyte infections with kinetics similar to the *sdhA* and *mavN* mutants in serum-dependent manner (Fig. 3c). Together these data are consistent with extracellular replication taking place in the absence of serum supplementation during infection. To test this idea with an alternative approach, we performed antibiotic protection assay using different spectinomycin concentrations to distinguish intracellular from extracellular replication. Spectinomycin is bacteriostatic and should impact extracellular replication at lower concentrations because host cell membranes limit drug diffusion. Indeed, spectinomycin at 30µg/ml completely blocked the replication of the *dotA* mutant in macrophage infections, which we posit to take place extracellularly, whereas the intracellular growth of the parental strain was only slightly reduced (SFig. 4a). Notably, both strains were equally sensitive to spectinomycin in axenic cultures (SFig. 4b and c) indicating that the distinct infection growth kinetics likely result from the inhibition of extracellular replication. In agreement, low dose spectinomycin (20µg/ml) completely blocked the growth of non-opsonized Lp in monocytes infections – condition in which the bacteria remain extracellular (SFig. 4d).

**Fig 3.**
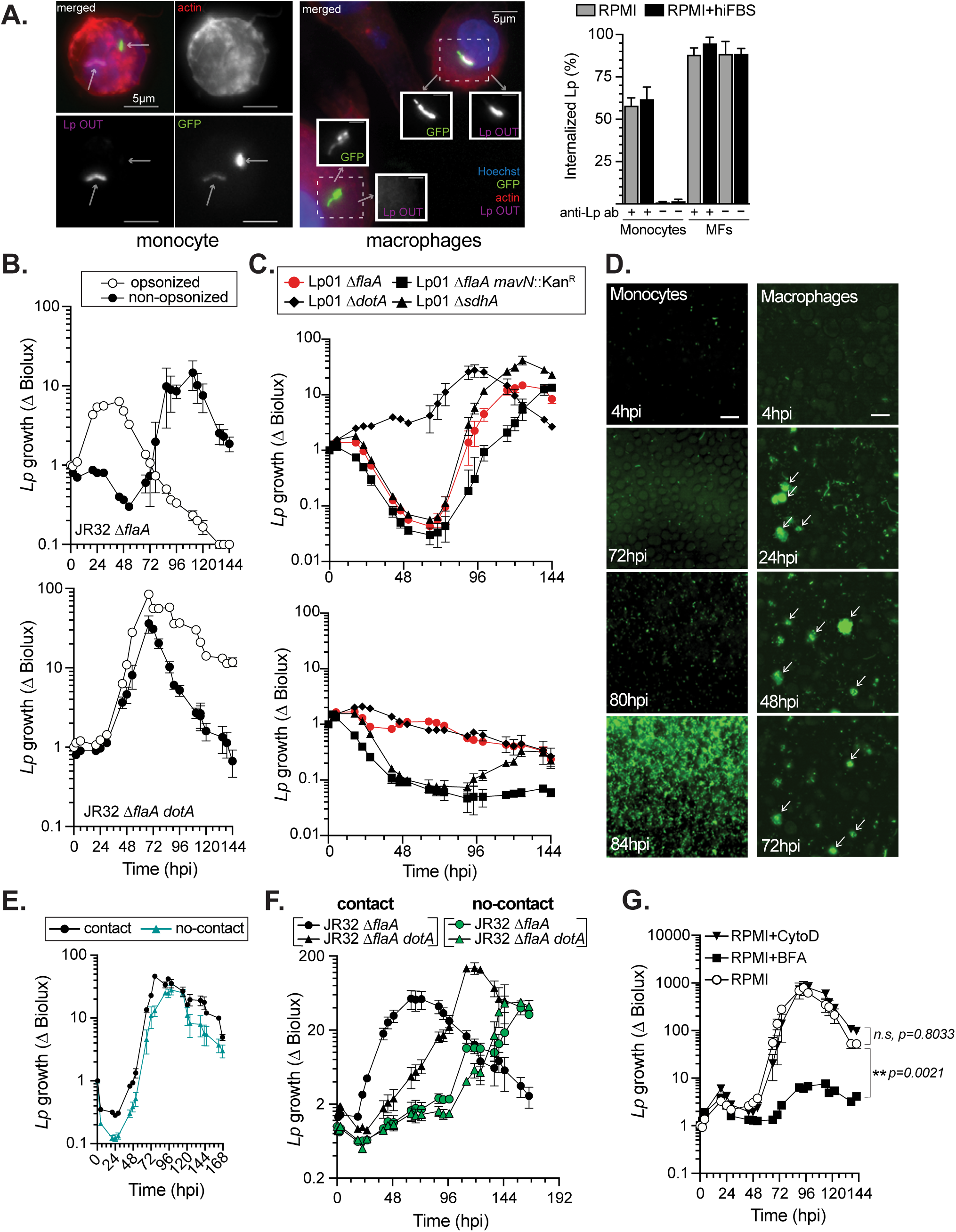
Monocytes and macrophages support extracellular Lp growth upon serum withdrawal. **(A)** Internalization of opsonized and non-opsonized Lp01 Δ*flaA* GFP+ by U937 cells at 6hpi. Representative z-projection micrographs from fixed cells strained by the inside/out method show both intracellular (single positive) and surface-associated (double positive) bacteria on monocytes (left panel) and macrophages (right panel). Graphed data represents averages of internalized bacteria for each condition from three technical replicates ± StDev (n > 150). Lp were opsonized with a polyclonal rabbit α-Lp and were stained with a chicken α-Lp IgY antibody. **(B)** Growth kinetics of the indicated Lp stains in cultures with U937 monocytes under SF conditions. Some bacteria were opsonized with a polyclonal rabbit α-Lp IgG for 30 min prior to infection. **(C)** Growth kinetics of the indicated Lp strains with U937 monocytes in the absence (top panel) or presence of hiFBS (bottom panel). **(D)** Representative micrographs from time-lapse live-cell microscopy showing growth kinetics of Lp01 Δ*flaA* GFP+ with U937 monocytes (left panels) and U937 macrophages (right panels). Source data is included in *Supplementary Video 1* and *2* (macrophages and monocytes respectively). Arrows denote individual LCVs in each frame. **(E-F)** Loss of direct bacterium-host cell contact alters Lp growth kinetics in macrophage but not monocyte cultures under SF conditions. **(E)** JR32 Δ*flaA* growth kinetics when cultured with U937 monocytes together or when separated by a 0.4µm membrane are comparable. **(F)** Loss-of-contact significantly delays initiation of JR32 Δ*flaA* replication in macrophage infections. **(G)** Lp01 Δ*flaA* growth kinetics with U937 monocytes in the presence of the actin-polymerization inhibitor cytochalasin D (5µM CytoD) or the secretory transport pathway inhibitor Brefeldin A (1μM BFA), which were added 60 min prior to infection. Statistical significance was determined by one-way ANOVA and the respective p-values are indicated. **(B-C, E-G)** Averages from three technical replicates ± StDev are shown. Two **(A, D)** or at least three **(B-C, E-G)** biological replicates were completed for each experiment.

Because the previous data is indicative of Lp extracellular replication in monocyte infections, we sought to visualize this process using live-cell imaging. To this end, monocytes or macrophages were cultured with GFP-expressing Lp in the absence of serum and were imaged every 4 hours over a period of several days spanning multiple rounds of infection (Fig. 3d and Svideo 1 and 2). As expected, Lp replication was spatially confined of the host cell resulting in the biogenesis of prototypical GFP+ spherical organelles that ultimately rupture releasing bacteria in the milieu (Fig. 3d and Svideo 1). Conversely, individual bacteria with unrestricted spatial lateral mobility were observed in monocyte cultures (Fig 3d. and Svideo 2), which indicates that Lp is not confined within the eukaryotic cell and is consistent with the uptake defect for non-opsonized Lp (Fig. 3a). Little bacterial replication was evident prior to ∼70hpi but the bacteria population increased afterwards dramatically within a short period of time (< 8 hours) (Fig. 3d and Svideo 2). Once bacterial replication started, the Lp GFP signal intensity increased uniformly throughout the imaging field in monocyte cultures, unlike the spatially localized signal produced in macrophage infections (Fig. 3d and Svideo 1 and 2). Thus, the live-cell imaging data clearly demonstrate the distinct properties of bacterial replication with host cells produced under conditions allowing or preventing bacterial uptake and strongly support the notion of Lp extracellular replication in specific infection settings.

Next, we investigated if Lp growth with monocytes required direct contact between the two cell types. Accordingly, infections were performed in a chamber assay, where bacteria are separated from the eukaryotic cells by a 0.4µm pore-seized membrane which allows only fluid phase exchange to occur between the two compartments. In this setting, Lp grew with similar kinetics regardless of whether direct contact with monocytes was allowed (Fig. 3e) confirming that the bacteria replicates extracellularly under those conditions. In the same setting, Lp growth was delayed when direct contact with macrophages was blocked by a physical membranous barrier (Fig. 3f) and phenocopied the contact-independent growth curve of the of the *dotA* mutant strain, which would be expected if both strains grew extracellularly. Furthermore, inhibition of actin polymerization with cytochalasin D resulting in a phagocytosis blockade did not interfere with Lp growth with monocytes (Fig. 3g). Based on the collective data we conclude that (i) Lp can replicate extracellularly in a contact-independent manner when cultured with monocytes or macrophages and (ii) serum presence restricts Lp extracellular growth.

### Macrophages and monocytes produce nutrients required for Lp extracellular replication

Lp is a fastidious bacterium with an amino acid-centric metabolism and is also an auxotroph for several amino acids which are acquired from the host cell during intracellular replication.^36,39^ Therefore, we sought to determine the nutrients source facilitating Lp extracellular growth. SF-RPMI by itself did not support bacterial growth (Fig. 4a), which would be expected because Lp cannot transport cystine across the outer membrane.^92^ Supplementation of RPMI with both cysteine and iron – another Lp growth factor^93^ – did not trigger bacterial replication either (Fig. 4a), indicating that the supplemented RPMI also lacks the capacity to support Lp growth. Because Lp can grow extracellularly in PRMI when macrophages or monocytes are present, one can posit that host cells likely produce one or more nutrients need by Lp for growth. Indeed, Lp grew in the absence of host cells when the bacteria were cultured in 0.2µm filtered conditioned medium (SF-RPMI), which was collected from 4 days-old macrophage cultures (Fig. 4a). Conditioned medium from unstimulated and *E.coli* LPS-stimulated macrophages supported Lp replication indistinguishably (Fig. 4a). Thus, we conclude that host cells are required and provide the necessary growth factors for Lp extracellular replication.

**Fig 4.**
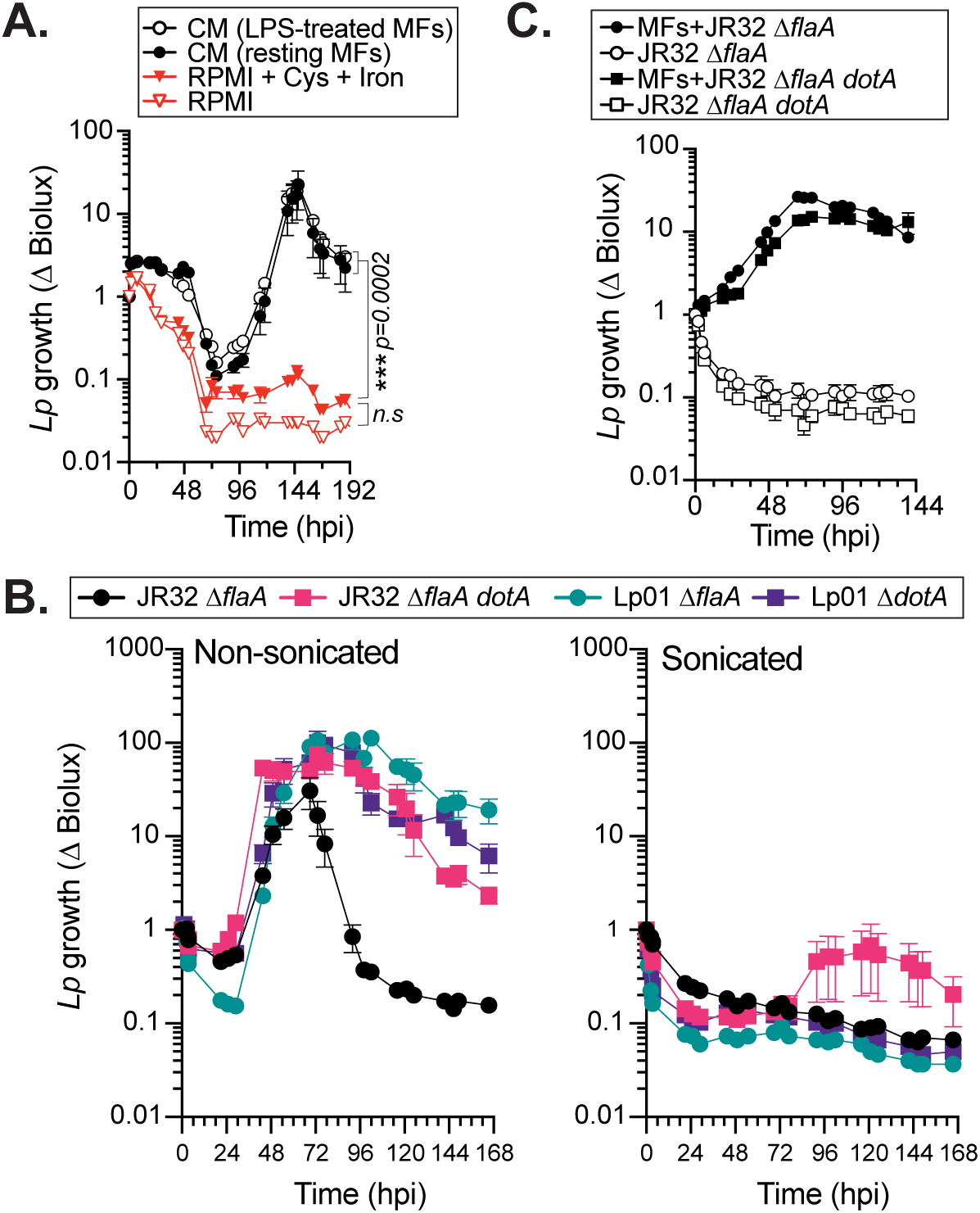
Nutrients secreted by live macrophages are sufficient and required for Lp extracellular replication. **(A)** Lp01Δ*flaA* growth kinetics in the absence of host cells. Bacteria were cultured in SF RPMI media supplemented with 3.3mM cysteine and 330µM Fe (NO_3_)_3_ or in 4 days RPMI conditioned media (CM) from resting or LPS-stimulated U937 macrophages (MFs). CMs were passaged through a 0.2µm membrane filter prior to bacteria addition. Statistical significance was determined by one-way ANOVA and the respective p-values are indicated. **(B)** Growth kinetics of the indicated Lp strains in the presence of live (non-sonicated) or physically ruptured (sonicated) U937 monocytes cultured in SF RPMI media. **(C)** Bacterial replication of the indicated strains in nutrient poor PBSG (PBS, 7.5mM glucose, 900µM CaCl_2_, 500µM MgCl_2_) media in the presence/absence of U937 macrophages. **(A-C)** Averages from three technical replicates ± StDev are shown. Representative of three biological replicates for each experiment is shown.

One possibility is that Lp grows extracellularly via necrotrophy fueled by nutrients released from dying macrophages/monocytes, which would explain the kinetic delay prior to initiation of extracellular replication as host cell viability gradually decreases. To investigate this scenario, Lp was cultured with monocytes that were sonicated in SF-RPMI prior to infection, which liberates intracellular host-derived metabolites for Lp consumption immediately (Fig. 4b). However, sonicated monocytes did not support the growth of either T4SS+ or T4SS-strains whereas the equivalent number of live (non-sonicated) monocytes did, demonstrating that at least some of the nutrients utilized by Lp during extracellular replication are produced by live host cells. To determine if macrophages can be the sole nutrients source for growth factors during Lp extracellular replication, akin to nutrients acquisition during intracellular replication, the growth assays were carried out in amino acids-free PBSG medium (PBS + 7.5mM glucose). Under those conditions, the *dotA* deletion mutant grew only in the presence of macrophages (Fig. 4c); thus, demonstrating that Lp acquires nutrients and other growth factors needed for extracellular growth from the host cells. Such a model would predict that nutrients concentrations sufficient for the initiation of bacterial replication would be reached faster in a smaller cell culture volume. Indeed, initiation of extracellular replication but not maximal growth positively correlated with the cell culture volume in monocytic infections, where bacterial replication started and peaked earlier when lower cell culture volumes were used during infection (SFig. 5 a-b). Altogether, we conclude that (i) macrophages/monocytes produce and release in the extracellular environment all nutrients necessary to support Lp replication and (ii) live host cells are required for at least the initiation of extracellular replication. In agreement, inhibition of the secretory pathway in monocytes with Brefeldin A (BFA) interfered with Lp extracellular replication (Fig. 3g). The exact nature of the host-derived nutrients and how they are sourced remains to be elucidated.

### Serum restrict Lp extracellular replication by scavenging bioavailable iron

It is apparent from our data that the practice of serum supplementation in cellular infection models restricted Lp extracellular replication, thus supporting the obligate intracellular growth paradigm in the field. To gain insight in the serum-mediated growth-restriction of Lp extracellular replication we focused on nutrients limitation as a plausible mechanism because heat-inactivated serum lacked microbiocidal activity towards Lp (SFig. 2b). Iron and cysteine are well-documented nutrient determinants of optimal Lp replication.^92,93^ Macrophages modulate the concentration of extracellular cysteine by direct export through the ASC neutral amino acid antiporters and by secretion of Thioredoxin, which reduces extracellular cystine to cysteine.^94,95^ Cysteine supplementation could not rescue Lp extracellular growth in macrophage infections when serum was present (Fig. 5a) indicating cysteine utilization is not limiting bacterial growth under those conditions. However, ferric iron supplementation fully restored Lp extracellular growth in the presence of either bovine (Fig. 5b-d) or human (Fig. 5e) serum in a dose-dependent manner (SFig. 6a) where the growth kinetics in the absence of serum were indistinguishable from the ‘serum + Fe^3+^’ condition (Fig. 5b-e). Supplementation of either ferrous or ferric iron abolished the serum-dependent restriction of Lp extracellular replication and triggered earlier bacterial replication under SF conditions (SFig. 6b), implicating iron availability in the extracellular milieu as a key growth determinant. The addition of other divalent metals (Zn^2+^, Mn^2+^ and Ni^2+^) did not counteract the serum restriction of Lp extracellular growth underlying the need specifically for iron (Fig. 5f). Conversely, iron supplementation minimally impacted Lp growth intracellularly (SFig. 6b), indicating bioavailable iron concentration inside macrophages is sufficient for optimal intracellular replication during infection. Importantly, primary human monocyte-derived macrophages (hMDMs) similar to U937 macrophages supported extracellular replication but only in the presence of exogenous iron when serum was present (Fig. 5e).

**Fig 5.**
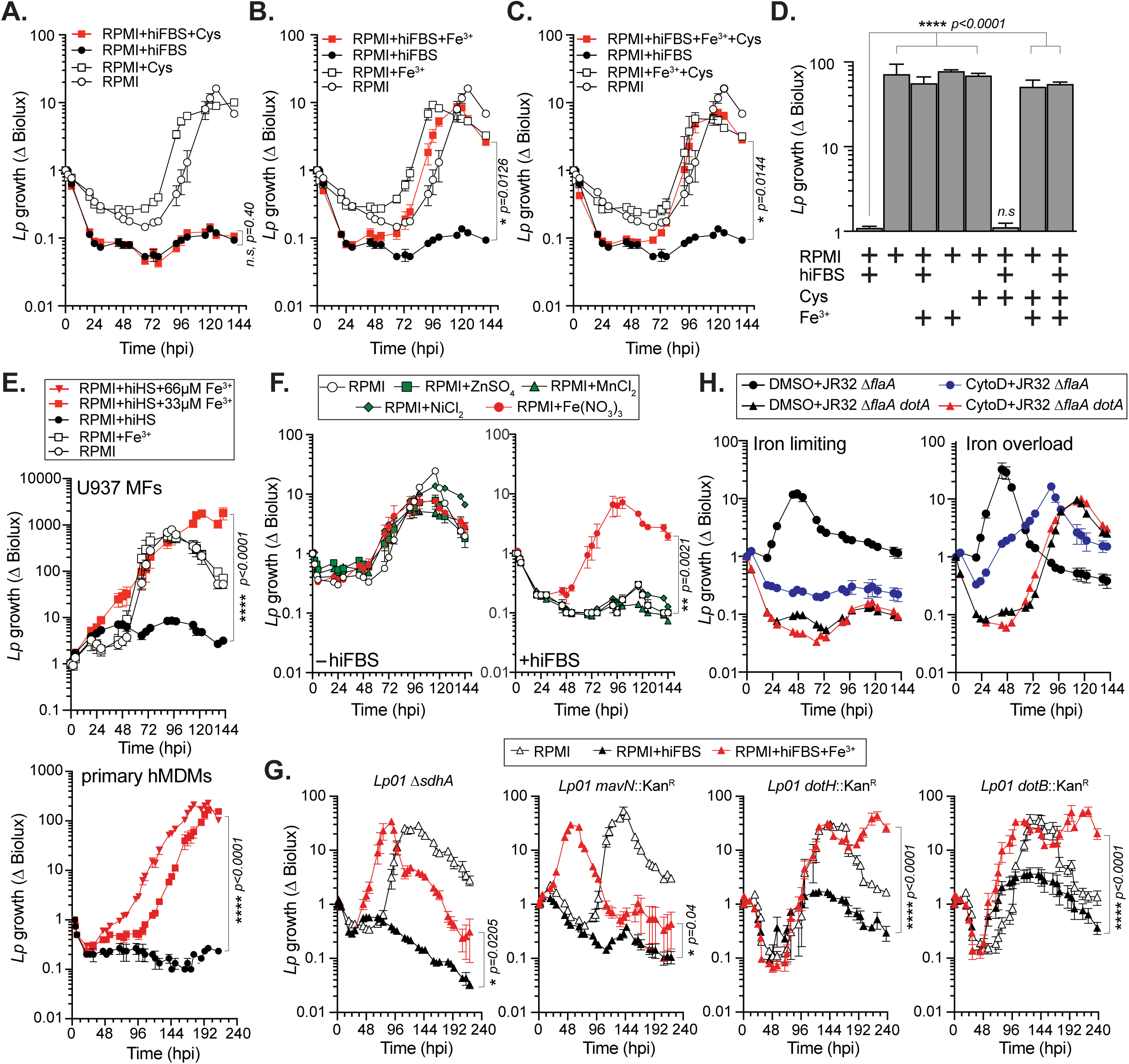
Iron overload overcomes serum restriction of *Legionella* extracellular growth. **(A-E)** Ferric iron but not cysteine rescues JR32 Δ*flaA dotA* growth with U937 macrophages **(A-E)** and primary human monocyte-derived macrophages (hMDMs) **(E)** in the presence of serum. Culture media supplementation with 33µM Fe(NO_3_)_3_ (Fe^3+^) **(B-E)** but not 3.3mM cysteine (Cys) **(A)** restores JR32 Δ*flaA dotA* growth in the presence of heat-inactivated FBS (hiFBS). **(D)** Fold difference in JR32 Δ*flaA dotA* bioluminescence output at 116 hpi normalized to the PRMI+hiFBS infection condition. **(E)** Culture media supplementation with the indicated Fe(NO_3_)_3_ (Fe^3+^) amounts rescues JR32 Δ*flaA dotA* growth in the presence of heat-inactivated human serum (hiHS). Note that hMDMs viability was rapidly lost in SF conditions and thus is not included in the analysis. **(F)** Non-iron divalent metal salts cannot restore JR32 Δ*flaA dotA* replication in the presence of hiFBS. **(G)** In the presence of serum, iron supplementation (33µM Fe(NO_3_)_3_) is sufficient to restore the growth of Lp mutants (*sdhA*, *mavN*, *dotH* and *dotB*) that cannot replicate intracellularly. **(H)** Ferric iron supplementation affects specifically Lp extracellular growth. Growth kinetics of the indicated Lp strains with U937 macrophages under *Iron limiting* (hiFBS) vs *Iron overload* (hiFBS+33µM Fe(NO_3_)_3_) conditions are shown. Bacterial internalization was blocked by the addition of 5µM cytochalasin D (CytoD) at 60 min prior to infection. **(A-H)** Averages from three technical replicates ± StDev are shown. Representative of three biological replicates for each experiment is shown. Statistical significance was determined by one-way ANOVA **(A-E, G)** and the respective p-values are indicated.

Iron-binding carrier proteins, such as Transferrin and Lactoferrin, are major serum constituents performing nutritional immunity functions by sequestering and limiting availability of divalent metals to bacterial pathogens.^67^ The iron-sequestration capacity of serum is a function of the saturation state of iron-binding proteins, thus iron supplementation can overload nutritional immunity defenses and restore iron availability for bacterial replication.^96,97^ Macrophages regulate extracellular iron levels, both locally and systemically, via dedicated uptake and secretion transport mechanisms.^98–100^ Without serum, the sole source of iron in our cell culture infection model remains the macrophage, which likely export sufficient amounts of iron in the extracellular milieu for Lp to growth. Therefore, we tested directly whether Lp extracellular replication specifically is dependent upon serum iron overload. To this end, the growth kinetics of different Lp mutant strains that cannot replicate intracellularly – *dotB*, *dotH*, *mavN* and *sdhA* – were measured under iron-limiting (i.e. ‘serum only’) or iron-overload (i.e. ‘serum+Fe^3+^’) conditions (Fig. 5g). For all strains, Lp extracellular growth was restored under iron-overload but not iron-limiting conditions (Fig. 5g). Notably, bacterial growth kinetics in iron-overload conditions either phenocopied (*dotH*) or were accelerated (*sdhA*, *mavN* and *dotH*) as compared to bacterial replication in the absence of serum (Fig. 5g). These data implicate serum iron-binding capacity as a critical regulator of Lp extracellular replication as demonstrated by strains that fail to replicate intracellularly. To assess serum capacity to block T4SS+ Lp extracellular growth supported by macrophages, we blocked phagocytosis with the actin-polymerization inhibitor cytochalasin D (cytoD) and measured bacterial replication under iron-overload and iron-limiting setting. Robust T4SS+ Lp replication was evident when iron-saturated serum and cytoD were present suggesting that macrophages can support extracellular replication of Dot/Icm+ Lp providing sufficient iron is available (Fig. 5h). Conversely, extracellular T4SS+ Lp replication was restricted under iron-limiting conditions (i.e. ‘+ serum and cyto D’), confirming the capacity of serum to restrict Lp extracellular replication specifically by limiting bioavailable iron released from macrophages (Fig. 5h). Similarly, the *dotA* deletion strain, which strictly replicates extracellularly, grew only under iron-overload conditions regardless of cytoD presence (Fig. 5h).

Because iron was critical for Lp extracellular growth in the presence of serum, we investigated if iron is required of Lp growth under SF conditions by treating cells with deferoxamine (DFO) – a non-toxic iron chelator that is clinically approved and effective for treatment of iron-overload pathological conditions.^101^ Treatment with 25µM DFO restricted Lp extracellular replication in monocyte infections as measured by a bioluminescence growth assay (Fig. 6a) and live-cell imaging (Fig. 6b). Importantly, addition of iron salt restored Lp growth in the presence of DFO demonstrating that the restriction of bacteria replication by DFO is due to iron-sequestration (Fig. 6a). Because iron is insufficient to trigger extracellular replication in SF conditions in the absence of macrophages (Fig. 4a), we conclude that iron is required but not sufficient for Lp extracellular replication during infection. Thus, as long as extracellular iron concentration does not exceed the sequestration capacity of nutritional immunity, macrophage/monocyte ability to support Lp extracellular replication is neutralized.

**Fig 6.**
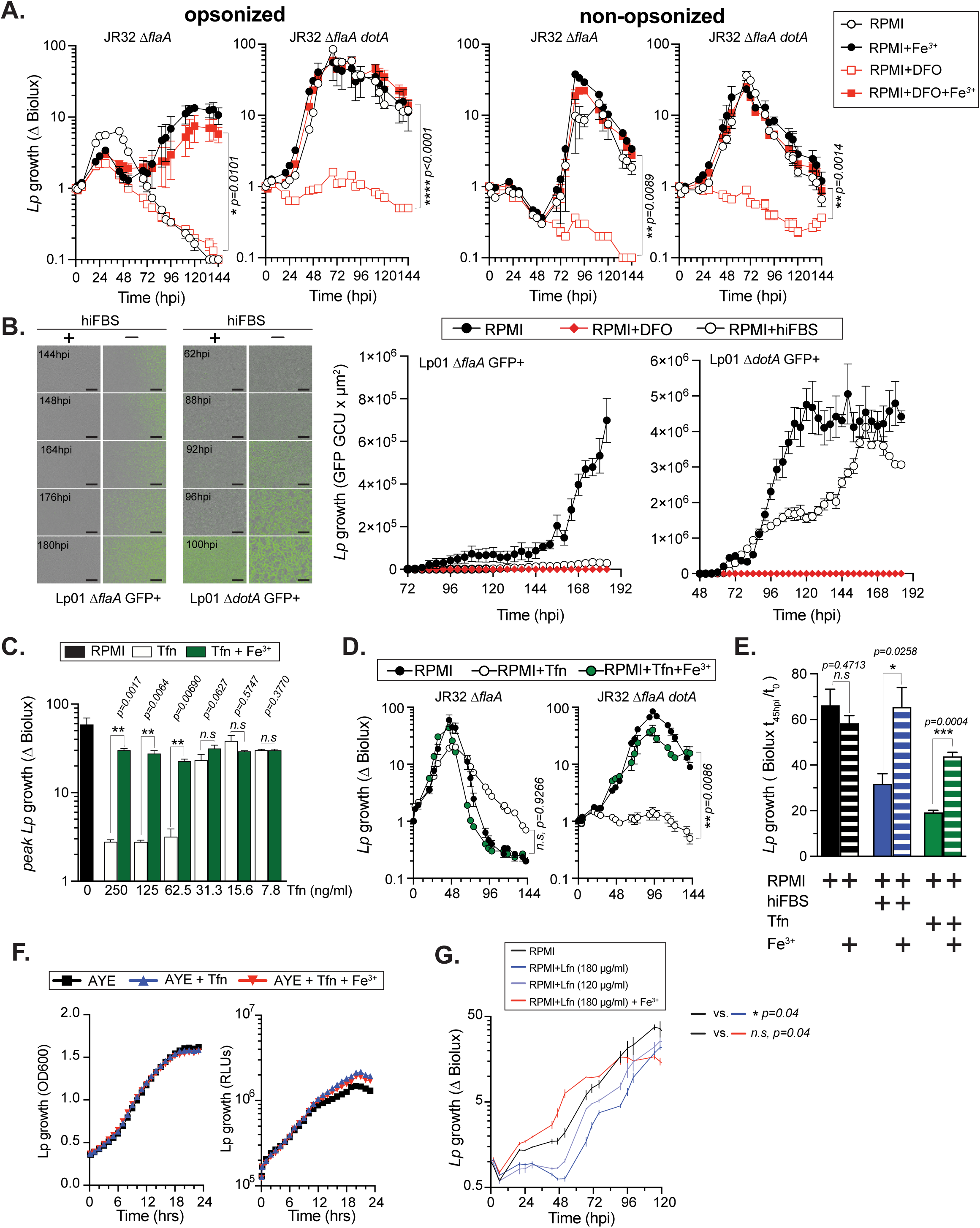
Sequestration of host-derived iron by nutritional immunity proteins restricts Lp extracellular replication. **(A)** The iron chelator Deferoxamine (DFO) blocks Lp extracellular growth in monocyte infections in an iron-dependent manner. Bacterial growth kinetics of the indicated Lp strains with U937 monocytes, where cells were treated with either 25µM DFO or 33µM Fe(NO_3_)_3_ individually or together. Growth kinetics of both opsonized and non-opsonized bacteria are shown. **(B)** Representative low magnification micrographs from time-lapse live-cell microscopy showing growth kinetics of non-opsonized Lp01 Δ*flaA* GFP+ and Lp01 Δ*flaA dotA* GFP+ with U937 monocytes in the presence of hiFBS or 25µM DFO. Bar = 100µm (left panels). Quantitative analysis of the changes in GFP signal total integrated intensity (GCU x µm^2^) over time in each well under the indicated growth conditions. **(C)** Supplementation of purified apo-Transferrin (Tfn) is sufficient to block Lp extracellular replication in a dose- and iron-dependent manner. Graph shows peak Lp01 Δ*dotA* growth (highest Biolux output / Biolux at T_0_) with U937 macrophages when Tfn was added at the indicated concentrations in the presence or absence of 330µM Fe(NO_3_)_3_. **(D)** Growth kinetics of the indicated Lp strains with U937 macrophages in SF RPMI under the indicated growth conditions (Tfn = 62.5µg/ml purified apo-Transferrin; Fe^3+^= 33µM Fe(NO_3_)_3_). **(E)** Maximum growth of JR32 Δ*flaA* in U937 macrophage infections under functional (+hiFBS or +Tfn) or compromised (+hiFBS/Fe^3+^ or +Tfn/Fe^3+^) nutritional immunity; Tfn = 62.5µg/ml purified apo-Transferrin, hiFBS = 10%v/v, Fe^3+^= 33µM Fe(NO_3_)_3_. **(F)** Apo-Transferrin does not reduce Lp growth or bioluminescence output *in vitro*. Lp growth in axenic liquid culture media AYE measured by the change in optical density (OD_600_) or Biolux output (RLUs) in the presence of Tfn and 330µM Fe(NO_3_)_3_. **(G)** Lactoferrin interferes with Lp extracellular growth in a dose-dependent manner. JR32 Δ*flaA dotA* growth kinetics with U937 macrophages under SF conditions in the presence of various concentration of purified human apo-Lactoferrin (Lfn). Fe^3+^ = 66µM Fe(NO_3_)_3_. **(A-G)** Averages from three technical replicates ± StDev are shown. Representative of two biological replicates for each experiment is shown. Statistical significance was determined by paired T-test **(C, E)** or one-way ANOVA **(A, D, G)** and the respective p-values are indicated.

### Host iron-binding proteins sustain nutritional immunity defenses against Lp

Transferrin and Lactoferrin are the major nutritional immunity iron-binding proteins in human airway secretions.^102^ Transferrin is a beta globulin with two high affinity ferric iron binding sites and the relative proportions of apo-, mono-, and diferric forms of Transferrin determines the percentage of Transferrin iron saturation in serum as well as its capacity to sequester additional iron molecules - a critical parameter for nutritional immunity. Because serum blocked Lp extracellular replication in an iron-dependent manner, we hypothesized that iron-binding carriers could mediate Lp restriction through iron-starvation. Therefore, the capacity of apo-Transferrin to restrict Lp extracellular growth in macrophage infections was investigated with the *dotA* mutant. The growth kinetics of the *dotA* mutant strain in macrophage cultures measured in the presence of different amounts of purified bovine apo-Transferrin (from 250.0µg/ml to 7.8µg/ml) clearly showed a dose-dependent restriction of Lp growth (Fig. 6c) that was reversed upon addition of ferric nitrate implying that Lp growth restriction is dependent on the iron-binding capacity of Transferrin (Fig. 6c). At 125µg/ml apo-Transferrin was microbiostatic against the *dotA* mutant in macrophage infections but also lowered the apex of the growth curve of the T4SS+ parental strain by ∼50 % without affecting the slope (Fig. 6d and e). Iron supplementation reversed the growth defect brought by apo-Transferrin under infection conditions. Importantly, apo-Transferrin did not reduce Lp viability nor bioluminescence output in axenic growth cultures (Fig. 6f). Lactoferrin also interfered with Lp extracellular replication in a manner that was abrogated by ferric iron (Fig. 6g). Thus, we conclude that iron-sequestration by host iron-binding proteins block Lp extracellular replication by absorbing iron released from macrophages during infection.

### Cooperation between cell intrinsic host defenses and nutritional immunity is required for restriction of Legionella growth

A robust pulmonary innate immune response orchestrated by TNF and IFNγ is critical for limiting Lp replication and resolving the infection by reprograming macrophages into a phenotypic state that is not permissive for intracellular bacterial replication.^52,53,103^ Primary human alveolar and monocyte-derived macrophages primed with IFNγ restrict Lp intracellular replication *in vitro*,^52,53^ emphasizing the importance IFNγ effector functions in Lp pathogenesis. Yet, the molecular determinants that allow a mild self-resolving legionellosis that is well-controlled by innate immunity and IFNγ to progress into Legionnaires’ disease - a severe atypical pneumonia – are unclear. Could extracellular replication driven by a pulmonary iron-overload potentially allow Lp to overcome innate immunity tilting the balance towards severe disease? It is well established that aging and history of cigarette smoking are major risk factors for severe legionellosis based on extensive epidemiological evidence.^5^ Smokers accumulate significantly higher concentration of iron in the alveolar compartment vs. non-smokers^104–109^ – a phenomenon that is exacerbated with age.^110^ Unlike iron, the amount of alveolar Transferrin remains unchanged in smokers^106–108^ indicating lowered pulmonary nutritional immune capacity in smokers may lead to iron-overload conditions sufficient to support Lp extracellular replication, which in turn would allow evasion of cell autonomous innate immune defenses, explosive bacterial growth and disease exacerbation. Such a scenario would be dependent on an intrinsic capacity of IFNγ stimulated macrophages to support Lp extracellular replication, especially under conditions of compromised nutritional immunity. Therefore, the capacity of macrophages to support Lp extracellular growth when primed with IFNγ alone or with IFNγ and *E.coli* LPS was investigated in the presence or absence of serum. First, we confirmed that IFNγ and IFNγ+LPS treatments elicited the appropriate transcriptional response in macrophages by demonstrating the induction of the interferon-stimulated genes *IDO1* and *IRF1*^111,112^ as well as the LPS regulated gene *IL6* (Fig. 7a). Next, the capacity of IFNγ-primed macrophages to restrict Lp intracellular replication was investigated via live-cell imaging. Macrophages primed with IFNγ in the presence of serum for 16hrs were infected with T4SS+ Lp expressing GFP in the presence of IFNγ under SF conditions and subsequently imaged for 24 hours (Fig. 7b). LCVs in non-primed macrophages increased in size unlike the LCVs in IFNγ-stimulated cells (Fig. 7b) indicating that serum withdrawal did not interfere with IFNγ-dependent restriction of Lp intracellular replication.

**Fig 7.**
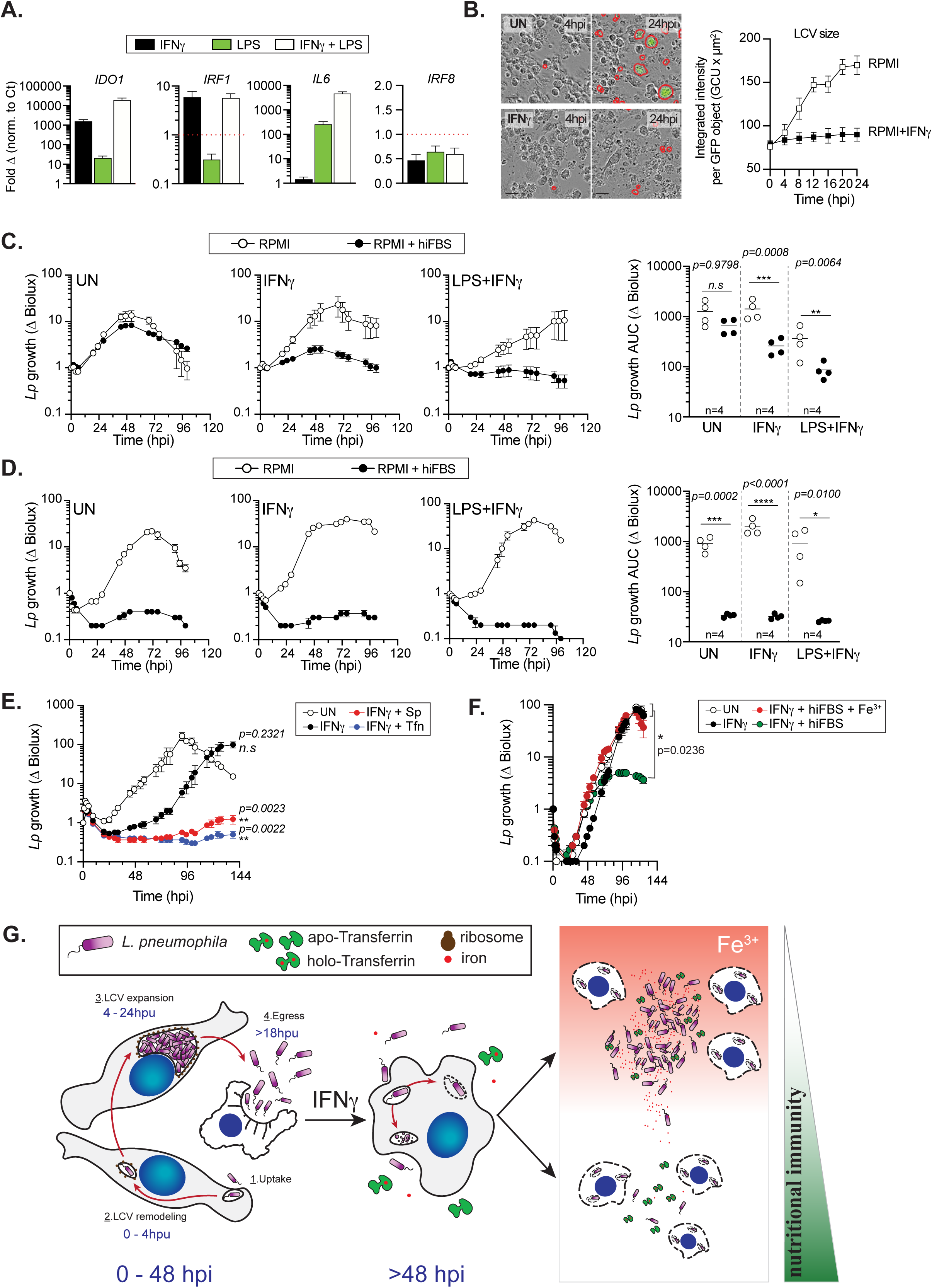
IFNγ and nutritional immunity cooperate to restrict Lp growth in macrophage infections. **(A)** qPCR analysis of IFNγ-inducible and inflammatory genes expression in U937 macrophages treated for 16hrs as indicated. For each gene expression data from each treatment is relative to respective GAPDH amount and is represented as fold change from unstimulated macrophages. **(B)** Quantitative analysis of LCVs expansion from time-lapse microscopy of U937 macrophages infected with Lp01 Δ*flaA* GFP+ for 24hrs that have been either primed with IFNγ (2µg/ml) for 16hrs or left unstimulated (UN). Serum was present during priming and was removed prior to the infection. Representative micrographs in which the LCVs are highlighted by a red border are shown in the left panel, whereas the change in LCV size (n > 200) is graphed in the right panel. **(C-D)** Lp growth restriction by macrophages requires both serum and IFNγ priming. Growth kinetics of JR32 Δ*flaA* **(C)** and JR32 Δ*flaA dotA* **(D)** with U937 macrophages primed with either IFNγ (2µg/ml) alone or IFNγ (2µg/ml) + *E.coli* LPS (100ng/ml) for 16hrs prior to infection in the presence of hiFBS (left panels). Area-under-the-curve analysis from four independent biological replicates are shown in the right panels. Treatments were maintained during the infections. In some conditions, the serum-containing RPMI was replaced with SF RPMI just before the inoculum was added. **(E)** Blockade of extracellular replication is required for the IFNγ-dependent restriction of Lp growth with human macrophages. Growth kinetics of JR32 Δ*flaA* in SF RPMI with unstimulated (UN) or IFNγ-primed U937 macrophages in the absence/presence of 20 µg/ml Spectinomycin (Sp) or 125 ng/ml purified apo-Transferrin (Tfn). IFNγ stimulation was maintained during the infections. **(G)** Proposed Lp pathogenesis model (see discussion). Averages from three technical replicates **(A-B, D-E)** or three biological replicates **(C)** ± StDev are shown. **(A-B, D-E)** Representative of three biological replicates for each experiment is shown. Statistical significance was determined by non-parametric Wilcoxon matched-pairs T-test **(C-D)** or one-way ANOVA **(E-F)** the respective p-values are indicated. n = number of biological replicates.

The growth of T4SS+ Lp with macrophages primed by either IFNγ or LPS+IFNγ was restricted in the presence of serum (Fig. 7c). However, when infections were carried out under SF conditions, T4SS+ Lp growth with IFNγ-primed macrophages was restored (Fig. 7c). Under the same conditions, the *dotA* mutant, which can only replicate extracellularly, grew with similar kinetics regardless of IFNγ priming, but strictly in the absence of serum (Fig. 7d). Thus, IFNγ-primed macrophages can support robust Lp extracellular replication. Considering that Lp intracellular growth is restricted by IFNγ in SF conditions (Fig. 7b), these data support a scenario where extracellular replication expands the bacterial population under conditions where IFNγ is present and nutritional immunity is absent. To test this scenario experimentally we utilized two distinct approaches to block Lp extracellular replication. The first was based on a low-dose spectinomycin antibiotic protection assay at concentrations that preferentially interfered with extracellular but not intracellular Lp replication (SFig 4) and the second was based on the capacity of apo-Transferrin to provide nutritional immunity and restrict Lp extracellular growth (Fig. 6d). In SF conditions, addition of either apo-Transferrin or low-dose spectinomycin was sufficient to restore the replication blockade imposed by IFNγ-primed macrophages (Fig. 7e), demonstrating that extracellular replication overcomes IFNγ restriction. Similarly, when nutritional immunity was compromised by iron-overload (serum + Fe^3+^) IFNγ-primed macrophages no longer restricted Lp growth (Fig. 7f). Previously published data with primary human cells demonstrating that iron supplementation overcomes IFNγ-mediated restriction of Lp replication also supports this notion,^76,77^ although the authors assumed that intracellular growth was restored. Notably, IFNγ- and LPS-priming significantly decreased expression of iron uptake and export regulators in macrophages indicating that iron release might potentially occur via a non-canonical mechanism independent of Ferroportin-1 (SFig. 7). Taken together, our data demonstrates that efficient host defenses to Lp require the joint action of IFNγ-driven cell mediated immunity and iron-sequestration by nutritional immunity proteins for the simultaneous blockade of intracellular and extracellular bacterial replication.

## Discussion

In the last four decades the idea that *Legionella* cannot replicate extracellularly under infection conditions has been a cornerstone of *Legionella* biology. This doctrine has shaped our view of *Legionella* pathogenicity in a way that parallels are drawn to obligate intracellular bacterial pathogens, where a ‘transmissive’ form exists outside of the host which transforms into a ‘replicative’ form within the intracellular niche, replicates and switches back to ‘transmissive’ form prior to egress.^51^ Our data challenges this longstanding notion by demonstrating that Lp readily replicates extracellularly during infection in the absence of or upon disruption of nutritional immunity. Serum utilization in tissue culture models of infections for decades has masked this phenomenon because of iron-dependent nutritional blockade. Several lines of evidence from our data demonstrate robust Lp extracellularly replication occurred when nutritional immunity is absent or is compromised due to saturation of iron-absorption capacity: (i) mutant strains with distinct intracellular growth defects now grew during infection; (ii) bacterial growth was observed even when host cells could not internalize bacteria; (iii) bacterial replication occurred despite a physical barrier separated Lp from the host cells; (iv) bacteria replicated in macrophage-conditioned medium in the absence of host cells altogether. We demonstrated that human immortalized (U937 monocytes and macrophages) as well as primary (hMDMs) cells support extracellular replication of *L. pneumophila* and *L. jordanis* when nutritional immunity is compromised. Thus, extracellular replication under infection conditions could be feature common to multiple pathogenic *Legionellaceae* species.

Notably, host cells were required for extracellular replication as macrophages and monocytes promoted bacterial growth likely by providing necessary nutrients missing from the cell culture media, which is evident by the conditioned medium experiments. We identified iron as one such milieu micronutrient that is required but is not sufficient for extracellular replication. Iron sequestration caused by serum-derived or purified iron-binding proteins imposed a blockade on extracellular replication that was abolished specifically under iron saturation conditions. Under SF conditions, the iron fueling Lp extracellular replication could only come from host cells either by Ferroportin mediated secretion or more likely by passive release from dying host cells because Ferroportin transcripts decreased by ∼80% in response to LPS in U937 macrophages (SFig 7). One iron-acquisition strategy evolved by human adapted bacterial pathogens to bypass nutritional immunity is the production of high-affinity bacterial receptors for Transferrin.^113–115^ However, apo-Transferrin interfered with Lp extracellular growth under infection conditions in a dose-dependent manner demonstrating that Transferrin blocks access to iron released from macrophages indicating that *Legionella* cannot source iron from holo-Transferrin directly and likely its siderophores have lower Fe^3+^ affinity compared to host iron-binding proteins.

Because legionellosis is caused by an environmental bacterium which co-evolved with unicellular amoebae and not a mammalian host, it has been difficult to explain how Lp escapes IFNγ-driven cell mediated immune defenses that efficiently block intracellular replication to cause severe disease. History of smoking as well as aging are the main risk factors for developing Legionnaires’ disease.^4,5^ Notably, bronchoalveolar lavage (BAL) fluid samples from smokers as compared to non-smokers are highly enriched in iron, without a concomitant increase in Transferrin.^104–109^ Similarly, aging has been shown to coincide with progressive accumulation of pulmonary iron,^110^ thus under both circumstances access to bioavailable iron is increased due to a state of lowered nutritional immunity. Indeed, BAL samples from smokers support higher growth of *Staphylococcus aureus* and *Pseudomonas aeruginosa* in an iron-dependent and Lactoferrin-dependent manner as compared to non-smokers.^116^ Moreover, a significant protective effect specifically against respiratory infections has been observed in the context of mild-to-moderate iron deficiency without an increase in cell-mediated immune responses.^72^ Could extracellular replication facilitated by iron overload potentially explain progression towards severe legionellosis? We demonstrated that iron-enriched milieu permits Lp replication outside of the host cell and bypass killing by IFNγ-primed macrophages. Previously, exogenous iron has been shown to rescue Lp growth with IFNγ-primed macrophages and the assumption was made that intracellular growth was restored.^76,77^ However, this assertion is incongruent with more recent data showing IFN stimulation results in (i) death of infected macrophages due to LCV rupture^57,58,60^ caused by the Guanylate binding protein 1 (GBP1) GTPase;^57^ (ii) downregulation of iron uptake via the Transferrin Receptor;^76^ (iii) an increase in iron export.^117^ Our work reconciles these observations by demonstrating that extracellular replication in iron-enriched milieus accounts for restoration of Lp growth with IFNγ-primed macrophages.

Another important aspect of our work is the discovery that outside of the host cell Lp can transition into a ‘replicative’ form under infection conditions by sourcing nutrients for extracellular growth from macrophages and monocytes. Lp cannot grow on glucose as the sole carbon source because of amino acid auxotrophy,^38^ yet our data revealed robust bacterial growth strictly in the presence of host cells in nutrient depleted medium lacking amino acids. Under such conditions, nutrients and growth factors can only be sourced from host cells. Even though heat-killed microorganisms (gram-negative bacteria as well as amoeba) have been shown to support limited Lp necrotrophic growth,^118^ in our assays sonicated monocytes did not support bacterial growth whereas live monocytes and macrophage-conditioned medium did. Thus, live monocytes/macrophages are necessary at least for initiation of extracellular bacterial replication during infection likely through production of critical nutrient(s) or a growth-initiating sensory cue; although, a potential role for necrotrophy cannot be excluded in the later growth stages. The sensory cues that trigger and the nutrients that fuel extracellular replication represent an important new aspect of Lp pathogenesis that warrants further research effort.

Our data argues for an updated pathogenesis paradigm taking into account nutritional immunity and extracellular bacterial replication. We propose the following model (Fig 7G). Upon inhalation and deposit in the lower respiratory tract, Lp likely undergoes several rounds of intracellular replication within alveolar macrophages triggering an inflammatory response, subsequent infiltration of innate immune cells, and localized IFNγ production by NK or NK T-cells. Once majority of bystander macrophages are primed by IFNγ, Lp intracellular growth would be restricted and the infection resolved providing nutritional immunity suppresses extracellular replication. This scenario is consistent with the self-resolving features of mild legionellosis in human infections (aka Pontiac fever) and low-dose infections in murine models.^4,5,119^ Because of the prolonged lag phage Lp extracellular replication likely coincides with IFNγ production and we argue that this is a critical junction in pathogenesis and disease progression. A defect in nutritional immunity caused by Transferrin saturation from a chronic pulmonary iron overload (as in patients with history of smoking) would establish an alveolar milieu permissive for extracellular replication further fueled by nutrients released by tissue-resident and infiltrating monocytes/macrophages. Under those conditions, extracellular replication can sustain bacterial growth despite IFNγ production further aggravating inflammation, eliciting the severe symptomatology associated with Legionnaires’ disease and causing pulmonary failure without a timely antibiotic administration.^120–122^ Such a model places an emphasis on extracellular survival and replication as important determinants of disease progression, assertion strongly supported by the dominance of clinical strains with the capacity to evade host defenses targeting extracellular bacteria. Majority of serogroup-1 clinical isolates unlike environmental isolates (80% vs 30%) express the *lag-1* gene encoding a LPS O-acetyltransferase, which confers resistance to complement killing and neutrophil internalization.^123^ Because ‘clinical’ strains are frequently isolated during outbreaks from symptomatic infections, the *lag-1* gene predominance in clinical isolates links extracellular antimicrobial defenses (i.e killing by complement and neutrophils) with clinical symptomatology and disease severity. Epidemiological data also argues for host-intrinsic rather than bacteria-intrinsic drivers of disease severity because outbreaks caused by a single clinical isolate frequently produces both mild and severe disease.^6–13^ Thus, extracellular replication as well as the defense mechanisms that counteract it are emerging as important previously underappreciated aspects of *Legionella* pathogenesis. Our work highlights the critical interdependency between nutritional and cell mediate immunity for generation of robust defense against bacterial infections and provides a reasonable mechanistic link between the well-documented deficiency in pulmonary nutritional immunity of smokers and severe legionellosis.

## Figure Legends

**SFig 1. Genetics and characterization of the *Legionella* strains encoding the *luxR* operon. (A)** Gene topology of the LuxR operon incorporated on the chromosome in the *L. pneumophila* (Lp) and *L. jordanis* (Lj) reporter strains used in this study. The *luxR* operon under the control of the *icmR* promoter was inserted on the chromosome via insertion duplication mutagenesis (*Lp icmRp*-*luxR*) for the generation of all bioluminescent reporter strains on the JR32 background used in the study. The *luxR* operon was inserted immediately downstream of the *icmR* gene on the chromosome via allelic exchange (*Lp icmR*-*luxR locus*) for the generation of all bioluminescent reporter strains on the Lp01 background used in this study. The *luxR* operon under the pTAC IPTG-inducible promoter together with the *lacI* repressor were incorporated within an intergenic region on the Lj chromosome via allelic exchange for the generation of all Lj bioluminescent reporter strains used in this study. **(B-D)** Axenic growth curves of *Legionella* strains encoding the indicated *luxR* reporters. The grey area curve indicates the change in optical density (OD_600_) over time. The dashed line indicates the changes in total bioluminescence of the bacterial culture over the indicated time period in the absence (blue) **(B-D)** or in the presence (red) **(D)** of IPTG. **(B-D)** Representative of three biological replicates for each experiment is shown.

**SFig 2. The effect of bovine and human serum on Lp replication in macrophage infections and axenic growth cultures.(A)** Growth kinetics of the indicated Lp strains with U937 macrophages in serum-free RPMI media (SFM) or RPMI supplemented with serum from different sources (10% v/v). Averages of three technical replicates are shown. One of two biological replicates is shown. **(B)** JR32 Δ*flaA dotA* growth kinetics in axenic AYE media or AYE supplemented with serum from different sources (10% v/v) as measured by changes in the culture optical density over time. Averages of three technical replicates ± StDev are shown. One of two biological replicates is shown. **(C)** Immunoblot analysis of the sera used in the experiments shown in (A) and (B). Transferrin band integrated intensity is shown for each lane on the blot and was normalized to the serum with the lowest Transferrin abundance. Total protein content for each serum was detected by Ponceau S staining as shown. **(D)** Total iron content of the sera used in the study as detected by ferrozine-based colorimetric assay. Averages of three technical replicates ± StDev are shown.

**SFig 3. L jordanis replication in macrophage infections and axenic growth cultures.**(**A)** Growth kinetics of the indicated Lj strains with U937 macrophages in serum-free RPMI media (SFM) or RPMI supplemented with bovine S4 serum (10% v/v). **(B)** Growth kinetics of the indicated Lj strains in axenic AYE media or AYE supplemented with bovine S4 serum (10% v/v) as measured by changes in the culture optical density over time. **(A-B)** Averages of three technical replicates ± StDev are shown. One of two biological replicates is shown for each experiment.

**SFig 4. Lp growth kinetics in spectinomycin treated macrophages and monocytes. (A-D)** Kinetics of Lp growth in the presence of spectinomycin (Sp). **(A and D)** Spectinomycin protection assay showing growth kinetics of the indicated Lp strains in U937 macrophages **(A)** and U937 monocytes **(D)** infections. **(B-C)** Axenic growth kinetics in AYE media of the indicated Lp strains in the absence or presence of 20 µg/ml spectinomycin as measured by changes in optical density (OD_600_) **(B)** or Biolux output (RLUs) **(C)**. **(A-D)** Averages of three technical replicates ± StDev are shown. One of three biological replicates is shown for each experiment.

**SFig 5. Lp growth kinetics dependency on the cell culture volume. (A-B)** Positive correlation between the cell culture volume and the kinetics of the peak Lp extracellular replication in U937 monocyte infections. Lp growth kinetics of the indicated strains in U937 monocytes tissue culture infections with different media volumes. Averages of three technical replicates ± StDev are shown. One of three biological replicates is shown for each experiment. **(B)** Correlational between cell culture volume during infection and (1) the time Lp growth peaks (left panel) and (2) the magnitude of Lp growth (right panel) in U937 monocyte infections for the indicated Lp strains. The Pearson *r* correlation coefficient (*r*) and *p* value is shown for each Lp strain. (two-tailed test, ** p < 0.004; ‘*n.s*’ – not significant)

**SFig 6. The effect of iron abundance on Lp growth with macrophages. (A)** Ferric Iron supplementation overcomes serum restriction of Lp extracellular replication in a dose-dependent manner. Growth kinetics of the indicated Lp strains with U937 macrophages in the presence of hiFBS and different concentrations of Fe(NO_3_)_3_. **(B)** Ferric and ferrous salts remove the nutritional blockade imposed by hiFBS. Growth kinetics of the indicated Lp strains with U937 macrophages in the presence/absence of hiFBS and different iron salts (330µM) – ferric nitrate, ferric sulfate and ferrous sulfate. **(A-B)** Averages of three technical replicates ± StDev are shown. One of three biological replicates is shown for each experiment.

**SFig 7. Effects IFNγ and LPS stimulation on expression of iron homeostasis regulatory genes. (A)** LPS and IFNγ stimulation reduced the expression of iron import and export regulatory genes in macrophages. qPCR analysis of U937 macrophages stimulated as indicated with either IFNγ (2µg/ml), *E.coli* LPS (100ng/ml) or both for 16hrs. Data for each gene is normalized to gene expression data from unstimulated macrophages. Averages of three biological replicates ± StDev are shown. *HAMP* – Hepcidin; *IREB2* - Iron Responsive Element Binding Protein 2; *TFCR* - Transferrin Receptor; *TFR2* - Transferrin Receptor 2; *FTH1* - Ferritin Heavy Chain 1; *SLC40A1* - Solute Carrier Family 40 Member 1 or Ferroportin-1.

**SVideo 1. Time-lapse live-cell imaging of Lp replication with U937 monocytes.** Growth kinetics of Lp01 Δ*flaA* GFP+ with U937 monocytes over an 84-hour period in serum-free RPMI (SFM) was imaged by time-lapse live-cell microscopy every 4 hours. Merged bright-field and fluorescence signals are shown at 1 frame per second. MOI = 2

**SVideo 2. Time-lapse live-cell imaging of Lp replication with U937 macrophages.** Growth kinetics of Lp01 Δ*flaA* GFP+ with U937 macrophages over an 84-hour period in serum-free RPMI (SFM) was imaged by time-lapse live-cell microscopy every 4 hours. Merged bright-field and fluorescence signals are shown at 1 frame per second. MOI = 2

## Materials and Methods

### Bacterial strains, plasmids, and media

The *L. pneumophila* strains used in this study were derived from the *L. pneumophila* serogroup 1, strain Lp01 or strain JR32^29^ and have a clean deletion of the *flaA* gene to avoid NLRC4-mediated pyroptotic cell death response triggered by flagellin. Importantly, strains lacking a functional T4SS as a result of inactivation of *dotA*, *dotB* or *dotH* do not trigger pyroptosis regardless of flagellin expression.

The bacterial strains, primers and plasmids used in this study are listed in Tables 1, 2 and 3 respectively. *Legionella* strains were cultured either on charcoal yeast extract (CYE) plates [1% yeast extract, 1%*N*-(2-acetamido)-2-aminoethanesulphonic acid (ACES; pH6.9), 3.3mM l-cysteine, 0.33mM Fe(NO_3_)_3_, 1.5% bacto-agar, 0.2% activated charcoal] or in complete ACES-buffered yeast extract (AYE) broth (10mg/ml ACES: pH6.9, 10mg/ml yeast extract, 400mg/l L-cysteine, 135mg/l Fe(NO_3_)_3_) supplemented with 100µg/ml streptomycin.^124^ As needed, the medium were supplemented with the following: 25 μg/ml kanamycin (Kam), 100 μg/ml streptomycin, 10 μg/ml chloramphenicol, 150mM sodium chloride, 1mM Isopropyl β-D-1-thiogalactopyranoside (IPTG), or different concentrations of spectinomycin.

**Table 1.** Strains used in this study.

| Strains | Genotype | Properties | Reference |
| --- | --- | --- | --- |
| CR19 | *E. coli* DH5α λpir *endA1 hsdR17 glnV44* (= *supE44*) *thi-1 recA1 gyrA96 relA1* ϕ80d*lac*Δ(*lacZ*)M15 $\Delta$(*lacZYA-argF*)U169 *zdg-232::Tn10 uidA::pir*<sup>+</sup> | conjugation helper strain | 125 |
| Lp01 Δ*flaA luxR* | *L. pneumophila* Lp01 Δ*flaA, icmR-luxR* locus | constitutive bioluminescence | This study |
| Lp01 Δ*dotA luxR* | *L. pneumophila* Lp01 Δ*dotA, icmR-luxR* locus | constitutive bioluminescence | This study |
| Lp01 Δ*sdhA luxR* | *L. pneumophila* Lp01 Δ*sdhA, icmR-luxR* locus | constitutive bioluminescence | This study |
| Lp01 Δ*flaA mavN::Kan<sup>R</sup> luxR* | *L. pneumophila* Lp01 Δ*flaA, mavN::Kan<sup>R</sup> icmR-luxR* locus | constitutive bioluminescence | This study |
| Lp01 *dotB::Kan<sup>R</sup> luxR* | *L. pneumophila* Lp01 *dotB::Kan<sup>R</sup> icmR-luxR* locus | constitutive bioluminescence | This study |
| Lp01 *dotH::Kan<sup>R</sup> luxR* | *L. pneumophila* Lp01 *dotH::Kan<sup>R</sup> icmR-luxR* locus | constitutive bioluminescence | This study |
| JR32 Δ*flaA LuxR* | *L. pneumophila* JR32 Δ*flaA, icmR<sub>P</sub>-luxR* | constitutive bioluminescence | This study |
| JR32 Δ*flaA* Δ*dotA LuxR* | *L. pneumophila* JR32 Δ*flaA* Δ*dotA, icmR<sub>P</sub>-luxR* | constitutive bioluminescence | This study |
| Lp01 Δ*flaA* GFP+ | *L. pneumophila* LP01 Δ*flaA* pPtac-GFP | inducible GFP expression, plasmid-borne pAM239 | 126 |
| Lp01 Δ*dotA* GFP+ | *L. pneumophila* LP01 Δ*dotA* pPtac-GFP | inducible GFP expression, plasmid-borne pAM239 | 126 |
| Lj *LuxR* | *L. jordanis* BL-540, *Strep<sup>R</sup>* Ptac-*luxR* | inducible bioluminescence, chromosome-borne | 80 |
| Lj Δ*dotB LuxR* | *L. jordanis* BL-540, *Strep<sup>R</sup>* Δ*dotB* Ptac-*luxR* | inducible bioluminescence, chromosome-borne | This study |

**Table 2.**
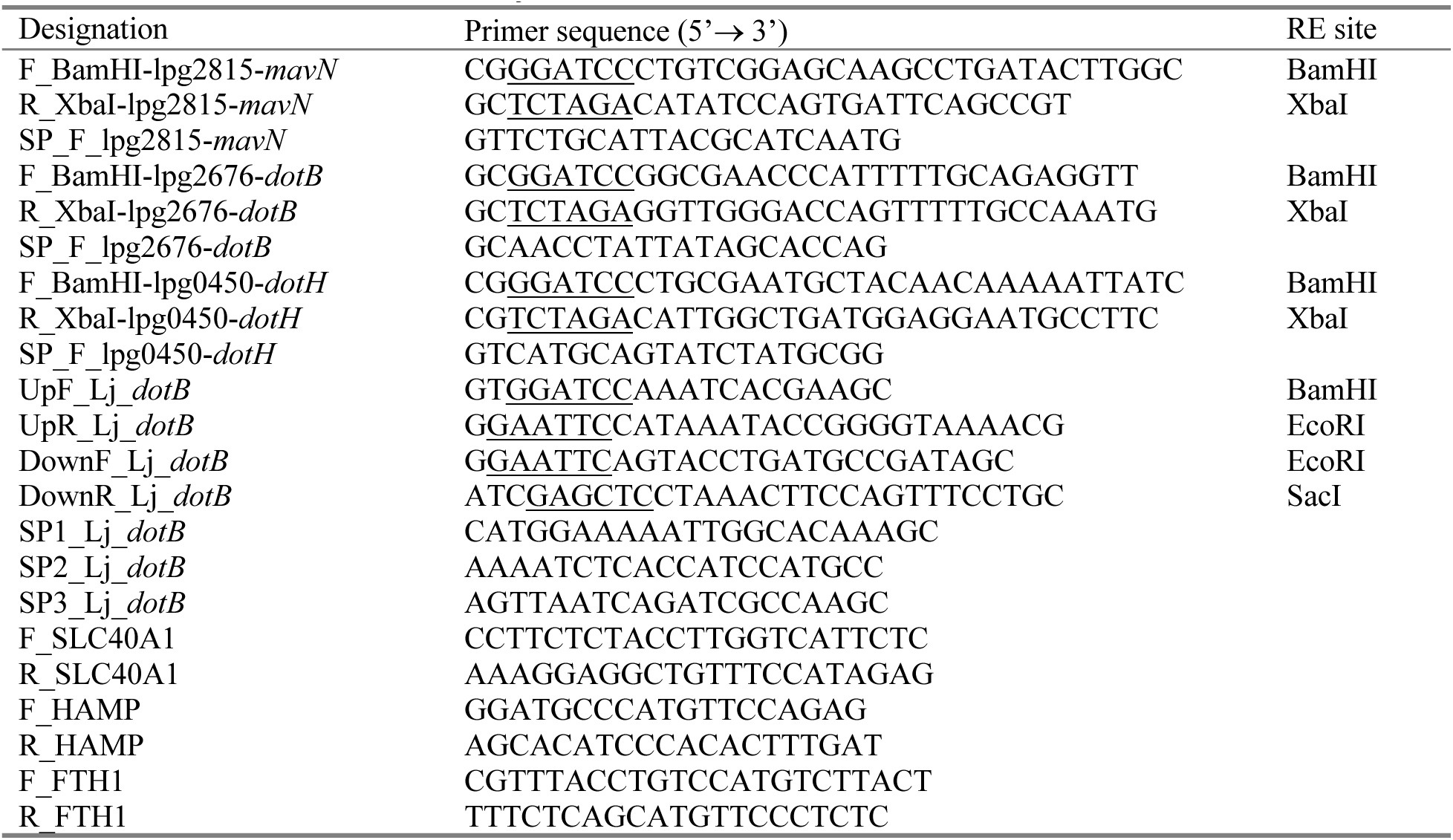
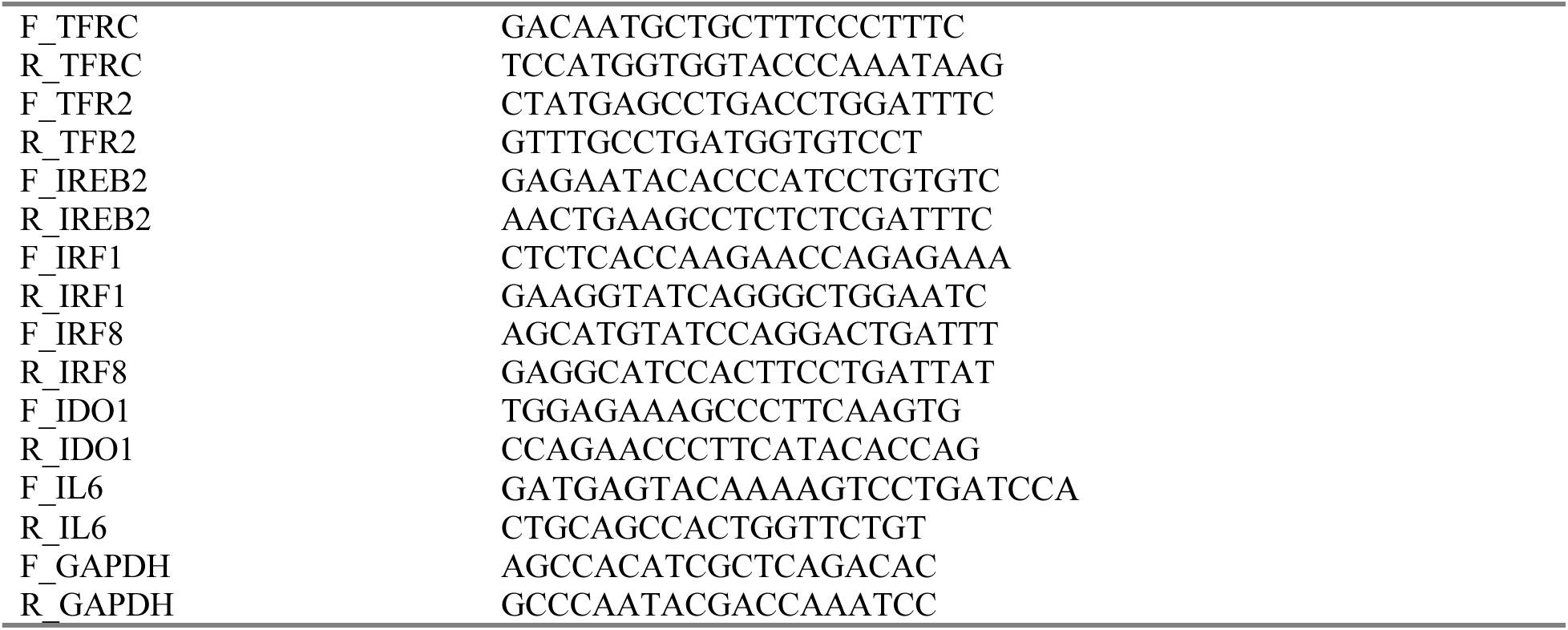
Primers used in this study.

**Table 3.** Plasmids used in this study.

| Plasmids | Important properties | Marker | Reference |
| --- | --- | --- | --- |
| pSR47 | *oriT* mobRP4 R6K *ori*, Kan<sup>R</sup> - suicide plasmid for insertion mutagenesis | Kan | 125 |
| pSR47s | pSR47, *sacB* Kan<sup>R</sup> - gene replacement vector | Kan | 125 |
| pSR47-icmRp-luxR | Plasmid encoding the *LuxR* operon under control of the Lp *icmR* promoter | Kan | 79 |
| pSR47s-icmR-luxR locus | Replacement chromosomal region containing the LuxR operon downstream of the Lp *icmR* gene | Kan | This study |
| pSR47-*mavN* | pSR47 containing a 995nt *mavN* intragenic region (137→1,131nt) | Kan | This study |
| pSR47-*dotB* | pSR47 containing a 786nt *dotB* intragenic region (151→936 nt) | Kan | This study |
| pSR47-*dotH* | pSR47 containing a 709nt *dotH* intragenic region (121→830 nt) | Kan | This study |
| pSR47s-CD *dotB<sub>Lj</sub>* | pSR47s containing the $\Delta dotB$ clean deletion allele | | This study |
| pSR47s-CD0376 | pSR47s containing the $\Delta sdhA$ clean deletion allele | | 126 |
| pSR47s-CD1340 | pSR47s containing the $\Delta flaA$ clean deletion allele | | 126 |

### Bacterial culture conditions for inoculum preparation

for all infection experiments, *Legionella* strains obtained from dense patches grown on CYE plates for two days were resuspended to OD_600_ of 0.5 U in 1ml AYE, placed in 15-ml glass culture tubes, and cultivated aerobically 24 to 26 hours with continuous shaking (175 rpm) at 37°C until early stationary phase was reached (OD_600_ range of 2.0 to 3.0 U). The *L. jordanis* reporter strains with the chromosomally encoded LuxR operon under the control of P_tac_ received 0.5mM IPTG at 8 hours post seeding in the AYE broth. The AYE broth cultures of *L. pneumophila* GFP+ strains harboring a plasmid carrying the *gfp* under the control of the P_tac_ contained 10μg/ml chloramphenicol and were supplemented with 0.1mM IPTG between 18 to 20 h post seeding in AYE broth.

### Generation of bioluminescent Legionella strains

all three bioluminescence producing regulons (see SFig 1A) were based on the *luxR* operon (*luxCDABE)* from *Photorhabdus luminescens*. The *L. pneumophila* JR32 strains used in the study encoded the constitutive *Lp icmRp-luxR* regulon, which has been previously described in detail.^79^ The *L. jordanis* strains encoded the IPTG-inducible *Lj pTAC-luxR* regulon, which has been previously described in detail.^80^ The *L. pneumophila* Lp01 strains used in the study encoded a new constitutive *Lp icmR-luxR locus* regulon. The *Lp icmR-luxR locus* is a synthetic regulon in which the *luxR* operon from *P. luminescens* is inserted immediately downstream of the Lp *icmR* gene in the context of the native *icmR* locus. For cloning, the synthetic *icmR* locus (1,943 nt) was obtained as a gBlock (IDT). It contained the Lp chromosome region spanning *icmT*, *icmS*, *icmR*, and *icmQ* genes and included four unique restriction enzyme sites: (i) *Bam*HI - localized upstream of the *icmT* gene; (ii) *Sac*I - localized downstream of the *icmQ* gene, as well as (iii) *Bgl*II and (iv) *Not*I localized in the intergenic region between the *icmR* and *icmQ* genes. The gBlock was digested with *Bam*HI/*Sac*I and cloned into similarly digested suicide plasmid pSR47s. Next, the 5,854nt *luxR* operon (*luxCDABE*) fragment was released from pXen13 with *Bam*HI and *Not*I, gel-purified and inserted between *icmR* and *icmQ* to yield pSR47s-icmR-luxR locus. The synthetic *icmR* regulon encoded by pSR47s-icmR-luxR locus was swapped with the native regulon in Lp01 strains by allelic exchange via double homologous recombination and was confirmed via bioluminescence emission; kanamycin sensitivity and sucrose resistance.^125^

### Inactivation of Lp genes via insertion mutagenesis

the single gene knockout strains lacking *mavN*, *dotB* or *dotH* were produced by intragenic insertion of the pSR47 plasmid on the Lp chromosome via homologous recombination. To this end, intragenic regions for *mavN* (137→ 1,131nt), *dotB* (151→ 937nt) and *dotH* (121→ 830nt) were PCR amplified from genomic DNA and cloned in pSR47 using primers listed in Table 2. The respective plasmids pSR47-*mavN*, pSR47-*dotB* and pSR47-*dotH* were introduced in Lp via tri-parental mating. Insertion mutants were isolated by kanamycin resistance and were genotyped by colony PCR to confirm correct insertion using the T3 promoter primer (5’-CAATTAACCCTCACTAAAGG-3’) together with the respective screening primer for each gene listed in Table 3.

### Inactivation of Lp genes via clean deletion (CD) mutagenesis

the *L. jordanis* Δ*dotB* allele was generated by fusing ∼1kb regions upstream and downstream of the *dotB* gene. To this end the upstream region was PCR amplified from genomic DNA using the primers UpF_Lj_*dotB* and UpR_Lj_*dotB*. The downstream region was similarly produced with the primers DownF_Lj_*dotB* and DownR_Lj_*dotB*. The resulting fragments were linked by an EcoRI site and cloned into the BamHI/SacI sites of the gene replacement vector pSR47s creating pSR47s-CD *dotB_Lj_*. The pSR47s-CD was electroporated (voltage-1800, capacitance- 25µF and resistance- 200ohms) in *L. jordanis* for the generation of the Δ*dotB* strain by allelic exchange of *dotB* with Δ*dotB* via double homologous recombination. The allelic exchange was confirmed by genotyping clonal isolates with the primers SP1_Lj_*dotB*, SP2_Lj_*dotB* SP3_Lj_*dotB* in a single PCR reaction that produces either a fragment of ∼300nt for *dotB* or a fragment of ∼450nt for Δ*dotB*. The clean deletion Lp strains lacking *flaA* or *sdhA* were generated with the gene replacement plasmids pSR47s-CD1340 and pSR47s-CD0376 via allelic exchange as detailed previously.^126^

### Reagents

Detailed reagents list is provided in Table 4. The purified polyclonal α-Lp IgY chicken antibody was custom generated by Cocalico Biologicals against formalin-killed bacteria.^126^ The purified polyclonal α-Lp IgG rabbit antibody (lot# Marsha) was a kind gift from Dr. Craig Roy (Yale University).

**Table 4.** Reagents used in this study.

| Reagent | Catalog # | Lot # | Supplier |
| --- | --- | --- | --- |
| Heat-inactivated Fetal Bovine Serum (FBS 1) | 512-048 | 16000-044 | Gibco |
| Heat-inactivated Fetal Bovine Serum (FBS 2) | 826-061 | 16000-044 | Gibco |
| Heat-inactivated Fetal Bovine Serum (FBS 3) | APJ 27610 | 30070.03 | Hyclone |
| Heat-inactivated Fetal Bovine Serum (FBS 4) | ABC 215874 | SH 30087.03 | Hyclone |
| Heat-inactivated Fetal Bovine Serum (FBS 5) | AC10335409 | SH 30910.03 | Hyclone |
| Heat-inactivated Fetal Bovine Serum (FBS 6) | 1983447 | 26140-079 | Gibco |
| Heat-inactivated Fetal Bovine Serum (FBS 7) | 30407 | S11150 | Atlanta Biologicals |
| Heat-inactivated Fetal Bovine Serum (FBS 8) | 30407 | E18055 | Atlanta Biologicals |
| Heat-inactivated Fetal Bovine Serum (FBS 9) | B21019 | S11150H | R&D Bovine Serum |
| Heat-inactivated Fetal Bovine Serum (FBS 10) | F-0500-D | F25E21D1 | Atlas Biologicals |
| Heat-inactivated Fetal Bovine Serum (FBS 11) | A 120030 | 900-208 | Gemeni Bioproducts |
| Heat-inactivated human serum (hS1) | 100-110 | H86W03J | Gemeni Bioproducts |
| Cytochalasin D | 11330 |  | Cayman Chemical |
| Deferoxamine | 14595 | 0618885-11 | Cayman Chemical |
| Purified Apo Transferrin from bovine plasma | J61626.MC | S071053 | Thermo Scientific |
| Human recombinant IFNγ | 570206 | B336905 | BioLegend |
| *E. coli* LPS | tlrl-eklps |  | Invivogen |
| RPMI-1640 | 25-506 | MS01GJ | Genesee |
| DMEM | 25-500 | MS0114L | Genesee |
| N-(2-acetamido)-2-aminoethanesulfonic acid | A11553-36 | 10239672 | Thermo Scientific |
| Yeast extract | 20-269 |  | APEX Bioreserach |
| Agar granulated | 20-248 | AB-2210138 | APEX Bioreserach |
| Activated charcoal | C9157-500G | SLBJ6388V | Sigma-Aldrich |
| Potassium hydroxide | 0489-1kg | 2746C296 | AMRESCO |
| RNeasy Mini kit (250) | 74106 |  | QIAGEN |
| DNaseI | M0303L |  | New England Biolabs |
| High-Capacity cDNA Reverse Transcription Kit | 4374966 |  | Thermo Fisher |
| Phorbol 12-myristic 13-acetate (PMA) | AG-CN2-0010-M001 |  | Adipogen |
| L-cysteine | A10435 | Q08H112 | Alfa Aesar |
| Fe(NO<sub>3</sub>)<sub>3</sub> | 33315 | W16A02 | Alfa Aesar |
| ZnSO<sub>4</sub>, | 19080 | N23069 | Research Organic |
| MnCl<sub>2</sub> | 155334 | 066J | MP Biomedicals |
| NiCl<sub>2</sub>, | N-5756 | 80K3489 | Sigma |
| Native Human Lactoferrin protein | AB285896 | 1004797-2 | ABCAN |
| Fe<sub>2</sub>SO<sub>4</sub>, | F1135 | 11KO171 | Sigma |
| FeSO<sub>4</sub> | F7002 | 50HO833 | Sigma |
| CuSO<sub>4</sub> | 0330-500G | 3586C030 | AMRESCO |
| Kanamycin | BP906-5 | 14446 | Fisher Scientific |
| Streptomycin | 8610 | 3551b38 | EM Science |
| Spectinomycin | 14324 | 056 1689-24 | Cayman Chemical |
| Chloramphenicol | 0230-100G | 1744CO11 | AMRESCO |
| Plasmid Mini Kit | 0230-100G | FH27305 | IBI Scientific |
| L-Ascorbic acid | 14656 | 0569615-46 | Cayman Chemical |
| Ferrozine | 32465 | 0648660-4 | Cayman Chemical |
| Neucoproine | 20874 | 0555221-5 | Cayman Chemical |
| Ammonium acetate | 09689 | 2111629 | Fluka Biochemika |
| Sodium hydroxide | 221465 | MKBS2033V | Sigma-Aldrich |
| Hydrochloric acid | HX0603-4 | 62262 | Sigma-Aldrich |
| Hoechst 33342 | H3570 |  | Life Technologies |
| Phalloidin-iFluor™ 594 Conjugate | 20553 |  | Cayman Chemical |
| goat α-chicken IgY-Alexa647 | A-21449 |  | Thermo Fisher |
| Monocyte Attachment Media | C-28051 |  | Sigma |

### Bacterial axenic growth assays

For axenic growth assays bacteria were collected from dense patches grown on CYE plates for two days, diluted in AYE with an initial OD_600_ = 0.5 U and were loaded onto a clear bottom white wall 96-well plate (Corning, cat# 3610). All experimental conditions were setup in triplicates and final culture volume was 200µl. The plate was loaded into Tecan Spark luminometer and incubated at 37°C for 24 h. Every 10 minutes, the cultures were agitated for 180 sec via a double orbital rotation at 108rpm and luminescence as well as OD_600_ data were collected. The bioluminescence output from each well was captured for 2 sec and quantified as a total relative light unit counts per second.

### Monocyte and macrophage cell culture conditions

Human U937 monocytes (ATCC, CRL1593.2) were cultured in RPMI-1640 with L-glutamine (BI Biologics, cat #01-100-1A), 10% v/v FBS and penicillin/streptomycin at a temperature of 37°C in the presence of 5% CO_2_. For macrophage differentiation, U937 monocytes were treated with 10 ng/ml Phorbol 12-myristate 13-acetate (PMA)(Adipogen) for the first 24 hours after which the cells were cultured for additional 48 hours without PMA and antibiotics. Human PBMCs were purchased from Sigma (cat#HUMANPBMC-0002644) and were differentiated into macrophage by continuous culture in RPMI-1640 with L-glutamine (BI Biologics, cat #01-100-1A), 20% v/v human serum (Sigma, cat# H4522) and penicillin/streptomycin at 37°C in the presence of 5% CO_2_ for 8 to 10 days, where the culture media were replaced every 4 days.

### Bioluminescence assay for *Legionella* replication with human monocytes and macrophages

For human U937 macrophages infections, U937 monocytes were seeded at a density of 1×10^5^ cells per well in white-wall clear-bottom 96-well plates (Corning cat# 3610) and differentiated into macrophages for 72 hours as indicated in “*Monocyte and macrophage cell culture conditions*”. For infection, U937 monocytes were seeded at the same cell density in 100µl of SF-RPMI or PBSG (PBS supplemented with 7.5mM Glucose, 0.9mM CaCl_2_, and 0.7mM MgCl_2_) per well for 3 hours prior to infection. The culture media used in each assay is indicated in the figure legends. Inoculum and other reagent were added to a final volume of 200µl/well. Primary human monocyte-derived macrophages (hMDMs) were seeded at 1×10^5^ cells per well in RPMI-1640 supplemented with human serum (10% v/v) (Sigma, cat# H4522) for 16 hours prior to infection. Because of rapid loss of viability infections with hMDMs were only carried out in the presence of human serum. Infections of U937 monocytes and macrophages were carried out under SF conditions or in the presence of FBS (10% v/v) as indicated for each assay with synchronized liquid culture grown bacteria at MOI=5. Unless indicated, the culture volume during infection was 200µl/well. Plates were centrifuged for 5 min at 500rpm after inoculum addition to enhance bacteria contact with the host cells. Plates were kept in a tissue culture incubator at 37°C with 5% CO_2_ and periodically, the bioluminescence output from each well was acquired by a Tecan Spark luminometer (integration time = 2 sec). The data is presented as fold change in bioluminescence from the T_0_ reading. All conditions in all assays were performed in technical triplicates.

### Infections of IFNγ-primed U937 macrophages

cells were cultured with either IFNγ (2µg/ml) or IFNγ + *E.coli* LPS (100ng/ml) for 16 hours in RPMI-1640 supplemented with hiFBS (10% v/v). At the time infection the culture medium was replaced with RPMI +/− hiFBS. The IFNγ and IFNγ + *E.coli* LPS treatments were maintained for the duration of the infection. The infections were carried out as outlined above and bacterial growth was measured via bioluminescence output.

### Infections with opsonized bacteria

as indicated in certain monocyte infections Lp was opsonized with a polyclonal rabbit anti-Lp IgG (1:1000 dilution) for 30 min prior to infection at room temperature in SF-RPMI. The infections were carried out as outlined above and bacterial growth was measured via bioluminescence output.

### Infections of sonicated monocytes

U937 monocytes were suspended in SF-RPMI at 2×10^6^ cells per 300µl. One group of cells was ruptured via sonication in Bioruptor^®^ Pico for 5 min (30sec ON, 30sec OFF sonication cycle) and another group was not sonicated. Volume equivalents containing 1×10^5^ cells (∼15µl) were added in each well from a 96 well plate to which the inoculums of liquid grown bacteria were added directly. The infections were carried out as outlined above and bacterial growth was measured via bioluminescence output. The final cell culture volume during infection was 200µl.

### Contact-dependent bacterial replication assay

U937 monocytes or macrophages were seeded in 24-well plates at 2.5×10^5^ cells/well. A transwell plate insert with a 0.4µm pore size bottom PET membrane (VWR, Cat# 76313-906) was placed in each well and the inoculum (MOI=5) was added either to the transwell insert for measurement of contact-independent bacterial replication or in the well to measure contact-dependent growth. The plate was centrifuged for 5min at 500rpm to bring the bacteria in contact with the host cell. Plates were kept in a tissue culture incubator at 37°C with 5% CO_2_ and periodically, the bioluminescence output from each well was acquired with a Tecan Spark luminometer (integration time = 2 sec). The data is presented as fold change in bioluminescence from the T_0_ reading. All conditions in all assays were performed in technical triplicates. Transwell membrane integrity in the contact-independent growth conditions was confirmed by plating serial dilutions of the contents from the lower chamber on CYE agar, and as expected viable CFU were not recovered.

### Bioluminescence assay for *Legionella* growth in conditioned media

For preparation of macrophage conditioned medium U937 monocytes were differentiated in 6 wells plates at 2×10^6^ cells/well and cultured with SF RPMI in the presence of absence of 100ng/ml *E.coli* LPS for 4 days. The conditioned medium was collected, centrifuged (10,000rpm, 5min) and passed through a 0.2µm pore filter. The collected media was either immediately used in bacterial growth assays or stored at -80°C for later use. Bacterial growth assay was setup in 96-well white wall plates and monitored by bioluminescence output, where 5×10^6^ bacteria were added to 200µl conditioned medium in each well. In control conditions, bacteria were added directly to SF RPMI in the presence or absence of 3.3mM cysteine + 330µM Fe (NO_3_)_3_. All, experimental conditions were carried out in technical triplicates.

### Colony Forming Units assay for *Legionella* growth

Macrophages were seeded, infected, and treated as listed in “*Bioluminescence assay for Legionella intracellular replication*”. Plates were kept in a tissue culture incubator at 37°C and 5% CO_2_ and periodically removed to collect bacteria. For bacterial recovery, cell culture medium from the well was moved into a 1.5ml Eppendorf tube and replaced with 100µl of sterile water for lysis of the infected macrophages via hypotonic shock and the plate was returned to the incubator for 10mins. Next, the contents of the well were pipetted at least 20 times and were collected in the respective Eppendorf tube which already contained the cell culture media. Eppendorf tubes were vortexed for 30sec and contents (∼300µl) were serially diluted five times for a total of six dilutions. 25µl from each dilution was plated on a CYE agar plate and incubated until colonies were visible. Colonies from a single dilution were counted and CFUs were calculated for each condition. Eight randomly selected colonies from each experimental condition at every time point were replicate streaked on CYA plates in the presence or absence of 150mM NaCl to confirm that all T4SS- strains used in the assay were Salt^R^, whereas T4SS+ strains were Salt^S^. All infection conditions for every experiment were performed in three technical replicates. The data is presented as fold change in recovered CFU over T_0_.

### Automated time-lapse live-cell imaging of *Legionella* growth with host cells

U937 monocytes or macrophages were seeded at 1×10^5^ in 96-well black-wall clear-bottom plates (Corning, cat# 3904) in phenol red-free DMEM supplemented as indicated for each assay. All infections were carried out at MOI = 2.5 in 150µl per well with AYE broth grown bacteria in the presence of 1mM IPTG. In each experiment, all conditions were performed in technical triplicates. Plates were centrifuged (500 rpm, 5min) and were loaded into the IncuCyte™ S3 microscope housing module. The IncuCyte™ S3 HD live-cell imaging platform (Sartorius) is a wide-filed microscope mounted inside a tissue culture incubator and is controlled by IncuCyte™ software. For each well, four single plane images in bright field and green channel (ExW 440-480nm/EmW 504-544nm) were automatically acquired with *S* Plan Fluor 20X/0.45 objective every four hours over several days. Images were analyzed with the IncuCyte™ Analysis software. For quantitation of bacterial replication, GFP signal was used for the generation of a binary mask that defined total signal integrated intensity (GCU x µm^2^) in each image or for each GFP+ object when LCV expansion was measured. The IncuCyte™ S3 imaging analysis was performed in the Innovative North Louisiana Experimental Therapeutics program (INLET) core facility at LSU Health-Shreveport.

### Gene expression analysis by quantitative real-time PCR

U937 macrophages were treated with 2μg/ml IFNγ, 100ng/ml *E.coli* LPS, or a combination of both for 16 hours, then total RNA was extracted with the RNeasy Mini kit (Qiagen), treated with DNase I (NEB, cat# 50-814-118) and cDNA was prepared with the High Capacity cDNA reverse transcriptase kit (Thermo Fisher, cat# 4368814). Real-time qPCR amplification of the cDNA was completed with PowerUp SYBR Green master mix (Thermo Fisher, cat# A25741), following the manufacturer’s instructions. For each reaction, 1 μl cDNA (equivalent to 0.1 - 0.2 μg of total RNA) and 20 pmoles of each gene-specific primer set (listed in Table 2) were combined in a 20 μl reaction volume. The LightCycler 96 Real-Time PCR System (Roche Diagnostics) was used for amplification with the following thermal cycle conditions: pre-amplification step at 95°C for 10min; amplification step of 40 cycles at 95°C for 10 sec, 60°C for 10 sec, and 72°C for 10 sec; followed by a melting step at 95°C for 10 sec, 65°C for 60 sec, and 97°C for 1 sec. For each experiment, samples were analyzed in triplicates. GAPDH expression served as the internal reference for normalization to determine the relative expression levels of specific transcripts, and fold differences were calculated using the 2^-ΔΔCt^ method.

### Bacteria uptake assay

To adhere U937 monocytes onto glass cover slips placed inside a 24-well plate, 1×10^5^ cells/well were cultured with Monocyte Attachment Medium (Sigma, cat# C-28051) for 2 hours after which the cells attached, and the medium was replaced with RPMI. U937 macrophages were directly differentiated onto glass cover slips at the same cell density. Infections were carried out with AYE-grown Lp01 Δ*flaA* GFP+ that were either opsonized or not opsonized with a polyclonal rabbit α-Lp IgG for 60 min prior to infection. All experimental conditions were done in technical triplicates for each experiment. After the bacteria were added to each well at MOI = 5, plates were centrifuged (500rpm, 5min) and the infection was allowed to proceed for 6 hours in the presence of 1mM IPTG and in the presence/absence of hiFBS (10% v/v). Next, cells were washed with warm PBS (3X), the infection was stopped by adding 2% paraformaldehyde (PFA) for 60 min at ambient temperature. The surface-associated bacteria were immunolabeled without plasma membrane permeabilization with chicken α-*Legionella* IgY antibody for 90 min in PBS containing goat serum (2% vol/vol) after which coverslips were washed with PBS (3X), fixed with 2% PFA (30 min) and permeabilized with 0.2% Triton X-100 (20min). After washing with PBS (3X), the samples were stained with Phalloidin-iFluor 594 Conjugate (1:2000), goat α-chicken IgY-Alexa647 (Thermo Fisher, cat# A-21449) (1:500 dilution) and with Hoechst (1:2000 dilution) for 60min in PBS containing goat serum (2% vol/vol). Coverslips were mounted with ProLong glass anti-fade mountant (ThermoFisher, cat# P36984) onto slides. Imaging was performed with inverted wide-field microscope (Nikon Eclipse Ti) controlled by NES Elements v4.3 imaging software (Nikon) using a 60X/1.40 oil objective (Nikon Plan Apo λ), LED illumination (Lumencor) and CoolSNAP MYO CCD camera. Image acquisition parameters - Hoechst (ExW 395 / EmW 455); GFP (ExW 470 / EmW 525), iFluor (ExW 555 / EmW 605) and Alexa647 (ExW 640 / EmW 705). The z-axis acquisition was set based on the out-of-focus boundaries and the distance between individual Z-slices was kept at 0.3µm. Only linear image corrections in brightness or contrast were completed. For each condition, over 150 bacteria were imaged and scored as either intracellular (single positive – green only) or not-internalized (double positive – green/red).

### Analysis of iron and Transferrin content in sera

The iron content of 12 commercially sourced sera from different manufacturers was determined by a ferrozine colorimetric assay. Ferrozine-based colorimetric assay. Briefly 100ml of serum was mixed with 100ml of 10mM HCl and 100ml of freshly prepared iron-releasing reagent (1.4M HCl, and 4.5% (w/v) KMnO_4_ in H_2_O). Next the samples were incubated for 2 hours at 60°C, cooled to ambient temperature and 30ml of the iron-detection reagent (6.6mM ferrozine, 6.5mM neocuproine, 2.5M ammonium acetate, and 1M ascorbic acid dissolved in water) was added. After 30 min incubation, 280µl of each sample was moved to one well of 96-well plate and the absorbance was read at 550nm on a Tecan Spark microplate reader. The iron content of each sample was calculated by comparing it absorbance to that of a range of concentration standards.

Immunoblot analysis for the Transferrin content in 2µl volume from each serum was performed with a rabbit polyclonal α-Transferrin IgG (ProteinTech, cat# 17435-1-AP) after standard SDS-PAGE. The total protein content transferred on the nitrocellulose membrane for each sample was detected after incubation with Ponceau S staining solution (Thermo Scientific, cat# A40000279) prior to immunoblotting.

### Statistical analysis

Calculations for statistical differences were completed with Prism v10.0.3 (GraphPad Software). The statistical tests applied for the different data sets are indicated in the figure legends and the resultant p-values are shown in the figures.

## Acknowledgments

This work was supported by grants from following funding sources: NIAID (AI143839) to SI, NIGMS (P20GM134974-5749) to AMD, Ike Muslow Predoctoral Fellowship from LSU Health Shreveport to AAW. We would like to thank the INLET High-Throughput Imaging Core at LSUHSC-Shreveport for technical assistance.

## References

1. Boamah, D.K., Zhou, G., Ensminger, A.W., and O’Connor, T.J. (2017). From Many Hosts, One Accidental Pathogen: The Diverse Protozoan Hosts of Legionella. Front Cell Infect Microbiol 7, 477. 10.3389/fcimb.2017.00477.

2. Gomez-Valero, L., and Buchrieser, C. (2013). Genome dynamics in Legionella: the basis of versatility and adaptation to intracellular replication. Cold Spring Harb Perspect Med 3. 10.1101/cshperspect.a009993.

3. Collier, S.A., Deng, L., Adam, E.A., Benedict, K.M., Beshearse, E.M., Blackstock, A.J., Bruce, B.B., Derado, G., Edens, C., Fullerton, K.E., et al. (2021). Estimate of Burden and Direct Healthcare Cost of Infectious Waterborne Disease in the United States. Emerg Infect Dis 27, 140–149. 10.3201/eid2701.190676.

4. Cunha, B.A., Burillo, A., and Bouza, E. (2016). Legionnaires’ disease. Lancet 387, 376–385. 10.1016/S0140-6736(15)60078-2.

5. Phin, N., Parry-Ford, F., Harrison, T., Stagg, H.R., Zhang, N., Kumar, K., Lortholary, O., Zumla, A., and Abubakar, I. (2014). Epidemiology and clinical management of Legionnaires’ disease. Lancet Infect Dis 14, 1011–1021. 10.1016/S1473-3099(14)70713-3.

6. Ambrose, J., Hampton, L.M., Fleming-Dutra, K.E., Marten, C., McClusky, C., Perry, C., Clemmons, N.A., McCormic, Z., Peik, S., Mancuso, J., et al. (2014). Large outbreak of Legionnaires’ disease and Pontiac fever at a military base. Epidemiol Infect 142, 2336–2346. 10.1017/S0950268813003440.

7. Benin, A.L., Benson, R.F., Arnold, K.E., Fiore, A.E., Cook, P.G., Williams, L.K., Fields, B., and Besser, R.E. (2002). An outbreak of travel-associated Legionnaires disease and Pontiac fever: the need for enhanced surveillance of travel-associated legionellosis in the United States. J Infect Dis 185, 237–243. 10.1086/338060.

8. Burnsed, L.J., Hicks, L.A., Smithee, L.M., Fields, B.S., Bradley, K.K., Pascoe, N., Richards, S.M., Mallonee, S., Littrell, L., Benson, R.F., et al. (2007). A large, travel-associated outbreak of legionellosis among hotel guests: utility of the urine antigen assay in confirming Pontiac fever. Clin Infect Dis 44, 222–228. 10.1086/510387.

9. 9. Euser, S.M., Pelgrim, M., and den Boer, J.W. (2010). Legionnaires’ disease and Pontiac fever after using a private outdoor whirlpool spa. Scand J Infect Dis 42, 910–916. 10.3109/00365548.2010.509331.

10. Girod, J.C., Reichman, R.C., Winn, W.C., Jr., Klaucke, D.N., Vogt, R.L., and Dolin, R. (1982). Pneumonic and nonpneumonic forms of legionellosis. The result of a common-source exposure to Legionella pneumophila. Arch Intern Med 142, 545–547.

11. Hunt, D.A., Cartwright, K.A., Smith, M.C., Middleton, J., Bartlett, C.L., Lee, J.V., Dennis, P.J., and Harper, D. (1991). An outbreak of Legionnaires’ disease in Gloucester. Epidemiol Infect 107, 133–141. 10.1017/s0950268800048767.

12. Smith, S.S., Ritger, K., Samala, U., Black, S.R., Okodua, M., Miller, L., Kozak-Muiznieks, N.A., Hicks, L.A., Steinheimer, C., Ewaidah, S., et al. (2015). Legionellosis Outbreak Associated With a Hotel Fountain. Open Forum Infect Dis 2, ofv164. 10.1093/ofid/ofv164.

13. Thomas, D.L., Mundy, L.M., and Tucker, P.C. (1993). Hot tub legionellosis. Legionnaires’ disease and Pontiac fever after a point-source exposure to Legionella pneumophila. Arch Intern Med 153, 2597–2599. 10.1001/archinte.153.22.2597.

14. Baskerville, A., Dowsett, A.B., Fitzgeorge, R.B., Hambleton, P., and Broster, M. (1983). Ultrastructure of pulmonary alveoli and macrophages in experimental Legionnaires’ disease. J Pathol 140, 77–90. 10.1002/path.1711400202.

15. Chandler, F.W., Blackmon, J.A., Hicklin, M.D., Cole, R.M., and Callaway, C.S. (1979). Ultrastructure of the agent of Legionnaires’ disease in the human lung. Am J Clin Pathol 71, 43–50. 10.1093/ajcp/71.1.43.

16. Davis, G.S., Winn, W.C., Jr., Gump, D.W., and Beaty, H.N. (1983). The kinetics of early inflammatory events during experimental pneumonia due to Legionella pneumophila in guinea pigs. J Infect Dis 148, 823–835. 10.1093/infdis/148.5.823.

17. Nash, T.W., Libby, D.M., and Horwitz, M.A. (1984). Interaction between the legionnaires’ disease bacterium (Legionella pneumophila) and human alveolar macrophages. Influence of antibody, lymphokines, and hydrocortisone. J Clin Invest 74, 771–782. 10.1172/JCI111493.

18. Winn, W.C., Jr., and Myerowitz, R.L. (1981). The pathology of the Legionella pneumonias. A review of 74 cases and the literature. Hum Pathol 12, 401–422. 10.1016/s0046-8177(81)80021-4.

19. Horwitz, M.A. (1983). Formation of a novel phagosome by the Legionnaires’ disease bacterium (Legionella pneumophila) in human monocytes. J Exp Med 158, 1319–1331. 10.1084/jem.158.4.1319.

20. Ivanov, S.S., and Roy, C.R. (2009). Modulation of ubiquitin dynamics and suppression of DALIS formation by the Legionella pneumophila Dot/Icm system. Cell Microbiol 11, 261–278. 10.1111/j.1462-5822.2008.01251.x.

21. Swanson, M.S., and Isberg, R.R. (1995). Association of Legionella pneumophila with the macrophage endoplasmic reticulum. Infect Immun 63, 3609–3620. 10.1128/iai.63.9.3609-3620.1995.

22. Tilney, L.G., Harb, O.S., Connelly, P.S., Robinson, C.G., and Roy, C.R. (2001). How the parasitic bacterium Legionella pneumophila modifies its phagosome and transforms it into rough ER: implications for conversion of plasma membrane to the ER membrane. J Cell Sci 114, 4637–4650. 10.1242/jcs.114.24.4637.

23. Cornejo, E., Schlaermann, P., and Mukherjee, S. (2017). How to rewire the host cell: A home improvement guide for intracellular bacteria. J Cell Biol 216, 3931–3948. 10.1083/jcb.201701095.

24. Personnic, N., Barlocher, K., Finsel, I., and Hilbi, H. (2016). Subversion of Retrograde Trafficking by Translocated Pathogen Effectors. Trends Microbiol 24, 450–462. 10.1016/j.tim.2016.02.003.

25. Roy, C.R., Berger, K.H., and Isberg, R.R. (1998). Legionella pneumophila DotA protein is required for early phagosome trafficking decisions that occur within minutes of bacterial uptake. Mol Microbiol 28, 663–674. 10.1046/j.1365-2958.1998.00841.x.

26. Elliott, J.A., and Winn, W.C., Jr. (1986). Treatment of alveolar macrophages with cytochalasin D inhibits uptake and subsequent growth of Legionella pneumophila. Infect Immun 51, 31–36. 10.1128/iai.51.1.31-36.1986.

27. Horwitz, M.A., and Silverstein, S.C. (1980). Legionnaires’ disease bacterium (Legionella pneumophila) multiples intracellularly in human monocytes. J Clin Invest 66, 441–450. 10.1172/JCI109874.

28. Andrews, H.L., Vogel, J.P., and Isberg, R.R. (1998). Identification of linked Legionella pneumophila genes essential for intracellular growth and evasion of the endocytic pathway. Infect Immun 66, 950–958. 10.1128/IAI.66.3.950-958.1998.

29. Berger, K.H., and Isberg, R.R. (1993). Two distinct defects in intracellular growth complemented by a single genetic locus in Legionella pneumophila. Mol Microbiol 7, 7–19. 10.1111/j.1365-2958.1993.tb01092.x.

30. Brand, B.C., Sadosky, A.B., and Shuman, H.A. (1994). The Legionella pneumophila icm locus: a set of genes required for intracellular multiplication in human macrophages. Mol Microbiol 14, 797–808. 10.1111/j.1365-2958.1994.tb01316.x.

31. Marra, A., Blander, S.J., Horwitz, M.A., and Shuman, H.A. (1992). Identification of a Legionella pneumophila locus required for intracellular multiplication in human macrophages. Proc Natl Acad Sci U S A 89, 9607–9611. 10.1073/pnas.89.20.9607.

32. Berger, K.H., Merriam, J.J., and Isberg, R.R. (1994). Altered intracellular targeting properties associated with mutations in the Legionella pneumophila dotA gene. Mol Microbiol 14, 809–822. 10.1111/j.1365-2958.1994.tb01317.x.

33. Segal, G., and Shuman, H.A. (1999). Legionella pneumophila utilizes the same genes to multiply within Acanthamoeba castellanii and human macrophages. Infect Immun 67, 2117–2124. 10.1128/IAI.67.5.2117-2124.1999.

34. Solomon, J.M., Rupper, A., Cardelli, J.A., and Isberg, R.R. (2000). Intracellular growth of Legionella pneumophila in Dictyostelium discoideum, a system for genetic analysis of host-pathogen interactions. Infect Immun 68, 2939–2947. 10.1128/IAI.68.5.2939-2947.2000.

35. Akinsulore, A., Owojuyigbe, A.M., Faponle, A.F., and Fatoye, F.O. (2015). Assessment of Preoperative and Postoperative Anxiety among Elective Major Surgery Patients in a Tertiary Hospital in Nigeria. Middle East J Anaesthesiol 23, 235–240.

36. Eisenreich, W., and Heuner, K. (2016). The life stage-specific pathometabolism of Legionella pneumophila. FEBS Lett 590, 3868–3886. 10.1002/1873-3468.12326.

37. George, J.R., Pine, L., Reeves, M.W., and Harrell, W.K. (1980). Amino acid requirements of Legionella pneumophila. J Clin Microbiol 11, 286–291. 10.1128/jcm.11.3.286-291.1980.

38. Pine, L., George, J.R., Reeves, M.W., and Harrell, W.K. (1979). Development of a chemically defined liquid medium for growth of Legionella pneumophila. J Clin Microbiol 9, 615–626. 10.1128/jcm.9.5.615-626.1979.

39. Tesh, M.J., and Miller, R.D. (1981). Amino acid requirements for Legionella pneumophila growth. J Clin Microbiol 13, 865–869. 10.1128/jcm.13.5.865-869.1981.

40. Horwitz, M.A. (1983). Cell-mediated immunity in Legionnaires’ disease. J Clin Invest 71, 1686–1697. 10.1172/jci110923.

41. Coleman, S.A., Fischer, E.R., Howe, D., Mead, D.J., and Heinzen, R.A. (2004). Temporal analysis of Coxiella burnetii morphological differentiation. J Bacteriol 186, 7344–7352. 10.1128/JB.186.21.7344-7352.2004.

42. Elwell, C., Mirrashidi, K., and Engel, J. (2016). Chlamydia cell biology and pathogenesis. Nat Rev Microbiol 14, 385–400. 10.1038/nrmicro.2016.30.

43. Gitsels, A., Sanders, N., and Vanrompay, D. (2019). Chlamydial Infection From Outside to Inside. Front Microbiol 10, 2329. 10.3389/fmicb.2019.02329.

44. Heinzen, R.A., Hackstadt, T., and Samuel, J.E. (1999). Developmental biology of Coxiella burnettii. Trends Microbiol 7, 149–154. 10.1016/s0966-842x(99)01475-4.

45. Troese, M.J., and Carlyon, J.A. (2009). Anaplasma phagocytophilum dense-cored organisms mediate cellular adherence through recognition of human P-selectin glycoprotein ligand 1. Infect Immun 77, 4018–4027. 10.1128/IAI.00527-09.

46. Zhang, J.Z., Popov, V.L., Gao, S., Walker, D.H., and Yu, X.J. (2007). The developmental cycle of Ehrlichia chaffeensis in vertebrate cells. Cell Microbiol 9, 610–618. 10.1111/j.1462-5822.2006.00812.x.

47. Bachman, M.A., and Swanson, M.S. (2004). Genetic evidence that Legionella pneumophila RpoS modulates expression of the transmission phenotype in both the exponential phase and the stationary phase. Infect Immun 72, 2468–2476. 10.1128/IAI.72.5.2468-2476.2004.

48. Byrne, B., and Swanson, M.S. (1998). Expression of Legionella pneumophila virulence traits in response to growth conditions. Infect Immun 66, 3029–3034. 10.1128/IAI.66.7.3029-3034.1998.

49. Garduno, R.A., Garduno, E., Hiltz, M., and Hoffman, P.S. (2002). Intracellular growth of Legionella pneumophila gives rise to a differentiated form dissimilar to stationary-phase forms. Infect Immun 70, 6273–6283. 10.1128/IAI.70.11.6273-6283.2002.

50. Hammer, B.K., Tateda, E.S., and Swanson, M.S. (2002). A two-component regulator induces the transmission phenotype of stationary-phase Legionella pneumophila. Mol Microbiol 44, 107–118. 10.1046/j.1365-2958.2002.02884.x.

51. Molofsky, A.B., and Swanson, M.S. (2004). Differentiate to thrive: lessons from the Legionella pneumophila life cycle. Mol Microbiol 53, 29–40. 10.1111/j.1365-2958.2004.04129.x.

52. Bhardwaj, N., Nash, T.W., and Horwitz, M.A. (1986). Interferon-gamma-activated human monocytes inhibit the intracellular multiplication of Legionella pneumophila. J Immunol 137, 2662–2669.

53. Nash, T.W., Libby, D.M., and Horwitz, M.A. (1988). IFN-gamma-activated human alveolar macrophages inhibit the intracellular multiplication of Legionella pneumophila. J Immunol 140, 3978–3981.

54. Price, J.V., Russo, D., Ji, D.X., Chavez, R.A., DiPeso, L., Lee, A.Y., Coers, J., and Vance, R.E. (2019). IRG1 and Inducible Nitric Oxide Synthase Act Redundantly with Other Interferon-Gamma-Induced Factors To Restrict Intracellular Replication of Legionella pneumophila. mBio 10. 10.1128/mBio.02629-19.

55. Shinozawa, Y., Matsumoto, T., Uchida, K., Tsujimoto, S., Iwakura, Y., and Yamaguchi, K. (2002). Role of interferon-gamma in inflammatory responses in murine respiratory infection with Legionella pneumophila. J Med Microbiol 51, 225–230. 10.1099/0022-1317-51-3-225.

56. Sporri, R., Joller, N., Albers, U., Hilbi, H., and Oxenius, A. (2006). MyD88-dependent IFN-gamma production by NK cells is key for control of Legionella pneumophila infection. J Immunol 176, 6162–6171. 10.4049/jimmunol.176.10.6162.

57. Bass, A.R., Egan, M.S., Alexander-Floyd, J., Lopes Fischer, N., Doerner, J., and Shin, S. (2023). Human GBP1 facilitates the rupture of the Legionella-containing vacuole and inflammasome activation. mBio 14, e0170723. 10.1128/mbio.01707-23.

58. Feeley, E.M., Pilla-Moffett, D.M., Zwack, E.E., Piro, A.S., Finethy, R., Kolb, J.P., Martinez, J., Brodsky, I.E., and Coers, J. (2017). Galectin-3 directs antimicrobial guanylate binding proteins to vacuoles furnished with bacterial secretion systems. Proc Natl Acad Sci U S A 114, E1698–E1706. 10.1073/pnas.1615771114.

59. Lippmann, J., Muller, H.C., Naujoks, J., Tabeling, C., Shin, S., Witzenrath, M., Hellwig, K., Kirschning, C.J., Taylor, G.A., Barchet, W., et al. (2011). Dissection of a type I interferon pathway in controlling bacterial intracellular infection in mice. Cell Microbiol 13, 1668–1682. 10.1111/j.1462-5822.2011.01646.x.

60. Liu, B.C., Sarhan, J., Panda, A., Muendlein, H.I., Ilyukha, V., Coers, J., Yamamoto, M., Isberg, R.R., and Poltorak, A. (2018). Constitutive Interferon Maintains GBP Expression Required for Release of Bacterial Components Upstream of Pyroptosis and Anti-DNA Responses. Cell Rep 24, 155–168 e155. 10.1016/j.celrep.2018.06.012.

61. Naujoks, J., Tabeling, C., Dill, B.D., Hoffmann, C., Brown, A.S., Kunze, M., Kempa, S., Peter, A., Mollenkopf, H.J., Dorhoi, A., et al. (2016). IFNs Modify the Proteome of Legionella-Containing Vacuoles and Restrict Infection Via IRG1-Derived Itaconic Acid. PLoS Pathog 12, e1005408. 10.1371/journal.ppat.1005408.

62. Pilla, D.M., Hagar, J.A., Haldar, A.K., Mason, A.K., Degrandi, D., Pfeffer, K., Ernst, R.K., Yamamoto, M., Miao, E.A., and Coers, J. (2014). Guanylate binding proteins promote caspase-11-dependent pyroptosis in response to cytoplasmic LPS. Proc Natl Acad Sci U S A 111, 6046–6051. 10.1073/pnas.1321700111.

63. Lettinga, K.D., Weijer, S., Speelman, P., Prins, J.M., Van Der Poll, T., and Verbon, A. (2003). Reduced interferon-gamma release in patients recovered from Legionnaires’ disease. Thorax 58, 63–67. 10.1136/thorax.58.1.63.

64. Murdoch, C.C., and Skaar, E.P. (2022). Nutritional immunity: the battle for nutrient metals at the host-pathogen interface. Nat Rev Microbiol 20, 657–670. 10.1038/s41579-022-00745-6.

65. Weinberg, E.D. (1974). Iron and susceptibility to infectious disease. Science 184, 952–956. 10.1126/science.184.4140.952.

66. Weinberg, E.D. (1975). Nutritional immunity. Host’s attempt to withold iron from microbial invaders. JAMA 231, 39–41. 10.1001/jama.231.1.39.

67. Neves, J., Haider, T., Gassmann, M., and Muckenthaler, M.U. (2019). Iron Homeostasis in the Lungs-A Balance between Health and Disease. Pharmaceuticals (Basel) 12. 10.3390/ph12010005.

68. Gangaidzo, I.T., Moyo, V.M., Mvundura, E., Aggrey, G., Murphree, N.L., Khumalo, H., Saungweme, T., Kasvosve, I., Gomo, Z.A., Rouault, T., et al. (2001). Association of pulmonary tuberculosis with increased dietary iron. J Infect Dis 184, 936–939. 10.1086/323203.

69. Ganz, T. (2009). Iron in innate immunity: starve the invaders. Curr Opin Immunol 21, 63–67. 10.1016/j.coi.2009.01.011.

70. Khan, F.A., Fisher, M.A., and Khakoo, R.A. (2007). Association of hemochromatosis with infectious diseases: expanding spectrum. Int J Infect Dis 11, 482–487. 10.1016/j.ijid.2007.04.007.

71. Sazawal, S., Black, R.E., Ramsan, M., Chwaya, H.M., Stoltzfus, R.J., Dutta, A., Dhingra, U., Kabole, I., Deb, S., Othman, M.K., and Kabole, F.M. (2006). Effects of routine prophylactic supplementation with iron and folic acid on admission to hospital and mortality in preschool children in a high malaria transmission setting: community-based, randomised, placebo-controlled trial. Lancet 367, 133–143. 10.1016/S0140-6736(06)67962-2.

72. Wander, K., Shell-Duncan, B., and Brindle, E. (2017). Lower incidence of respiratory infections among iron-deficient children in Kilimanjaro, Tanzania. Evol Med Public Health 2017, 109–119. 10.1093/emph/eox010.

73. Donovan, A., Brownlie, A., Zhou, Y., Shepard, J., Pratt, S.J., Moynihan, J., Paw, B.H., Drejer, A., Barut, B., Zapata, A., et al. (2000). Positional cloning of zebrafish ferroportin1 identifies a conserved vertebrate iron exporter. Nature 403, 776–781. 10.1038/35001596.

74. Donovan, A., Lima, C.A., Pinkus, J.L., Pinkus, G.S., Zon, L.I., Robine, S., and Andrews, N.C. (2005). The iron exporter ferroportin/Slc40a1 is essential for iron homeostasis. Cell Metab 1, 191–200. 10.1016/j.cmet.2005.01.003.

75. Cassat, J.E., and Skaar, E.P. (2013). Iron in infection and immunity. Cell Host Microbe 13, 509–519. 10.1016/j.chom.2013.04.010.

76. Byrd, T.F., and Horwitz, M.A. (1989). Interferon gamma-activated human monocytes downregulate transferrin receptors and inhibit the intracellular multiplication of Legionella pneumophila by limiting the availability of iron. J Clin Invest 83, 1457–1465. 10.1172/JCI114038.

77. Byrd, T.F., and Horwitz, M.A. (1991). Lactoferrin inhibits or promotes Legionella pneumophila intracellular multiplication in nonactivated and interferon gamma-activated human monocytes depending upon its degree of iron saturation. Iron-lactoferrin and nonphysiologic iron chelates reverse monocyte activation against Legionella pneumophila. J Clin Invest 88, 1103–1112. 10.1172/JCI115409.

78. Ren, T., Zamboni, D.S., Roy, C.R., Dietrich, W.F., and Vance, R.E. (2006). Flagellin-deficient Legionella mutants evade caspase-1-and Naip5-mediated macrophage immunity. PLoS Pathog 2, e18. 10.1371/journal.ppat.0020018.

79. Ondari, E., Wilkins, A., Latimer, B., Dragoi, A.M., and Ivanov, S.S. (2023). Cellular cholesterol licenses Legionella pneumophila intracellular replication in macrophages. Microb Cell 10, 1–17. 10.15698/mic2023.01.789.

80. Wilkins, A.A., Schwarz, B., Torres-Escobar, A., Castore, R., Landry, L., Latimer, B., Bohrnsen, E., Bosio, C.M., Dragoi, A.M., and Ivanov, S.S. (2023). The intracellular growth of the vacuolar pathogen Legionella pneumophila is dependent on the acyl chain composition of host membranes. bioRxiv. 10.1101/2023.11.19.567753.

81. Vogel, J.P., Roy, C., and Isberg, R.R. (1996). Use of salt to isolate Legionella pneumophila mutants unable to replicate in macrophages. Ann N Y Acad Sci 797, 271–272. 10.1111/j.1749-6632.1996.tb52975.x.

82. Kubori, T., Koike, M., Bui, X.T., Higaki, S., Aizawa, S., and Nagai, H. (2014). Native structure of a type IV secretion system core complex essential for Legionella pathogenesis. Proc Natl Acad Sci U S A 111, 11804–11809. 10.1073/pnas.1404506111.

83. Vincent, C.D., Friedman, J.R., Jeong, K.C., Buford, E.C., Miller, J.L., and Vogel, J.P. (2006). Identification of the core transmembrane complex of the Legionella Dot/Icm type IV secretion system. Mol Microbiol 62, 1278–1291. 10.1111/j.1365-2958.2006.05446.x.

84. Chetrit, D., Hu, B., Christie, P.J., Roy, C.R., and Liu, J. (2018). A unique cytoplasmic ATPase complex defines the Legionella pneumophila type IV secretion channel. Nat Microbiol 3, 678–686. 10.1038/s41564-018-0165-z.

85. Sexton, J.A., Pinkner, J.S., Roth, R., Heuser, J.E., Hultgren, S.J., and Vogel, J.P. (2004). The Legionella pneumophila PilT homologue DotB exhibits ATPase activity that is critical for intracellular growth. J Bacteriol 186, 1658–1666. 10.1128/JB.186.6.1658-1666.2004.

86. Izu, K., Yoshida, S., Miyamoto, H., Chang, B., Ogawa, M., Yamamoto, H., Goto, Y., and Taniguchi, H. (1999). Grouping of 20 reference strains of Legionella species by the growth ability within mouse and guinea pig macrophages. FEMS Immunol Med Microbiol 26, 61–68. 10.1111/j.1574-695X.1999.tb01372.x.

87. Creasey, E.A., and Isberg, R.R. (2012). The protein SdhA maintains the integrity of the Legionella-containing vacuole. Proc Natl Acad Sci U S A 109, 3481–3486. 10.1073/pnas.1121286109.

88. Isaac, D.T., Laguna, R.K., Valtz, N., and Isberg, R.R. (2015). MavN is a Legionella pneumophila vacuole-associated protein required for efficient iron acquisition during intracellular growth. Proc Natl Acad Sci U S A 112, E5208–5217. 10.1073/pnas.1511389112.

89. Choi, W.Y., Kim, S., Aurass, P., Huo, W., Creasey, E.A., Edwards, M., Lowe, M., and Isberg, R.R. (2021). SdhA blocks disruption of the Legionella-containing vacuole by hijacking the OCRL phosphatase. Cell Rep 37, 109894. 10.1016/j.celrep.2021.109894.

90. Abeyrathna, S.S., Abeyrathna, N.S., Thai, N.K., Sarkar, P., D’Arcy, S., and Meloni, G. (2019). IroT/MavN Is a Legionella Transmembrane Fe(II) Transporter: Metal Selectivity and Translocation Kinetics Revealed by in Vitro Real-Time Transport. Biochemistry 58, 4337–4342. 10.1021/acs.biochem.9b00658.

91. Christenson, E.T., Isaac, D.T., Yoshida, K., Lipo, E., Kim, J.S., Ghirlando, R., Isberg, R.R., and Banerjee, A. (2019). The iron-regulated vacuolar Legionella pneumophila MavN protein is a transition-metal transporter. Proc Natl Acad Sci U S A 116, 17775–17785. 10.1073/pnas.1902806116.

92. Ewann, F., and Hoffman, P.S. (2006). Cysteine metabolism in Legionella pneumophila: characterization of an L-cystine-utilizing mutant. Appl Environ Microbiol 72, 3993–4000. 10.1128/AEM.00684-06.

93. Reeves, M.W., Pine, L., Hutner, S.H., George, J.R., and Harrell, W.K. (1981). Metal requirements of Legionella pneumophila. J Clin Microbiol 13, 688–695. 10.1128/jcm.13.4.688-695.1981.

94. Castellani, P., Angelini, G., Delfino, L., Matucci, A., and Rubartelli, A. (2008). The thiol redox state of lymphoid organs is modified by immunization: role of different immune cell populations. Eur J Immunol 38, 2419–2425. 10.1002/eji.200838439.

95. Gmunder, H., Eck, H.P., Benninghoff, B., Roth, S., and Droge, W. (1990). Macrophages regulate intracellular glutathione levels of lymphocytes. Evidence for an immunoregulatory role of cysteine. Cell Immunol 129, 32–46. 10.1016/0008-8749(90)90184-s.

96. Bullen, J.J., Rogers, H.J., Spalding, P.B., and Ward, C.G. (2006). Natural resistance, iron and infection: a challenge for clinical medicine. J Med Microbiol 55, 251–258. 10.1099/jmm.0.46386-0.

97. Hunter, R.L., Bennett, B., Towns, M., and Vogler, W.R. (1984). Transferrin in disease II: defects in the regulation of transferrin saturation with iron contribute to susceptibility to infection. Am J Clin Pathol 81, 748–753. 10.1093/ajcp/81.6.748.

98. Ganz, T., and Nemeth, E. (2015). Iron homeostasis in host defence and inflammation. Nat Rev Immunol 15, 500–510. 10.1038/nri3863.

99. Nairz, M., Theurl, I., Swirski, F.K., and Weiss, G. (2017). “Pumping iron”-how macrophages handle iron at the systemic, microenvironmental, and cellular levels. Pflugers Arch 469, 397–418. 10.1007/s00424-017-1944-8.

100. Winn, N.C., Volk, K.M., and Hasty, A.H. (2020). Regulation of tissue iron homeostasis: the macrophage “ferrostat”. JCI Insight 5. 10.1172/jci.insight.132964.

101. Parker, J.B., Griffin, M.F., Downer, M.A., Akras, D., Berry, C.E., Cotterell, A.C., Gurtner, G.C., Longaker, M.T., and Wan, D.C. (2023). Chelating the valley of death: Deferoxamine’s path from bench to wound clinic. Front Med (Lausanne) 10, 1015711. 10.3389/fmed.2023.1015711.

102. Szabo, S., Barbu, Z., Lakatos, L., Laszlo, I., and Szabo, A. (1980). Local production of proteins in normal human bronchial secretion. Respiration 39, 172–178. 10.1159/000194213.

103. Horwitz, M.A., and Silverstein, S.C. (1981). Activated human monocytes inhibit the intracellular multiplication of Legionnaires’ disease bacteria. J Exp Med 154, 1618–1635. 10.1084/jem.154.5.1618.

104. Corhay, J.L., Weber, G., Bury, T., Mariz, S., Roelandts, I., and Radermecker, M.F. (1992). Iron content in human alveolar macrophages. Eur Respir J 5, 804–809.

105. Ghio, A.J., Hilborn, E.D., Stonehuerner, J.G., Dailey, L.A., Carter, J.D., Richards, J.H., Crissman, K.M., Foronjy, R.F., Uyeminami, D.L., and Pinkerton, K.E. (2008). Particulate matter in cigarette smoke alters iron homeostasis to produce a biological effect. Am J Respir Crit Care Med 178, 1130–1138. 10.1164/rccm.200802-334OC.

106. McGowan, S.E., and Henley, S.A. (1988). Iron and ferritin contents and distribution in human alveolar macrophages. J Lab Clin Med 111, 611–617.

107. Thompson, A.B., Bohling, T., Heires, A., Linder, J., and Rennard, S.I. (1991). Lower respiratory tract iron burden is increased in association with cigarette smoking. J Lab Clin Med 117, 493–499.

108. Wesselius, L.J., Flowers, C.H., and Skikne, B.S. (1992). Alveolar macrophage content of isoferritins and transferrin. Comparison of nonsmokers and smokers with and without chronic airflow obstruction. Am Rev Respir Dis 145, 311–316. 10.1164/ajrccm/145.2_Pt_1.311.

109. Wesselius, L.J., Nelson, M.E., and Skikne, B.S. (1994). Increased release of ferritin and iron by iron-loaded alveolar macrophages in cigarette smokers. Am J Respir Crit Care Med 150, 690–695. 10.1164/ajrccm.150.3.8087339.

110. Ghio, A.J., Pritchard, R.J., Dittrich, K.L., and Samet, J.M. (1997). Non-heme (Fe3+) in the lung increases with age in both humans and rats. J Lab Clin Med 129, 53–61. 10.1016/s0022-2143(97)90161-x.

111. Carlin, J.M., Borden, E.C., Sondel, P.M., and Byrne, G.I. (1989). Interferon-induced indoleamine 2,3-dioxygenase activity in human mononuclear phagocytes. J Leukoc Biol 45, 29–34. 10.1002/jlb.45.1.29.

112. Der, S.D., Zhou, A., Williams, B.R., and Silverman, R.H. (1998). Identification of genes differentially regulated by interferon alpha, beta, or gamma using oligonucleotide arrays. Proc Natl Acad Sci U S A 95, 15623–15628. 10.1073/pnas.95.26.15623.

113. Loosmore, S.M., Yang, Y.P., Coleman, D.C., Shortreed, J.M., England, D.M., Harkness, R.E., Chong, P.S., and Klein, M.H. (1996). Cloning and expression of the Haemophilus influenzae transferrin receptor genes. Mol Microbiol 19, 575–586. 10.1046/j.1365-2958.1996.406943.x.

114. Modun, B., Kendall, D., and Williams, P. (1994). Staphylococci express a receptor for human transferrin: identification of a 42-kilodalton cell wall transferrin-binding protein. Infect Immun 62, 3850–3858. 10.1128/iai.62.9.3850-3858.1994.

115. Noinaj, N., Easley, N.C., Oke, M., Mizuno, N., Gumbart, J., Boura, E., Steere, A.N., Zak, O., Aisen, P., Tajkhorshid, E., et al. (2012). Structural basis for iron piracy by pathogenic Neisseria. Nature 483, 53–58. 10.1038/nature10823.

116. Vargas Buonfiglio, L.G., Borcherding, J.A., Frommelt, M., Parker, G.J., Duchman, B., Vanegas Calderon, O.G., Fernandez-Ruiz, R., Noriega, J.E., Stone, E.A., Gerke, A.K., et al. (2018). Airway surface liquid from smokers promotes bacterial growth and biofilm formation via iron-lactoferrin imbalance. Respir Res 19, 42. 10.1186/s12931-018-0743-x.

117. Abreu, R., Essler, L., Giri, P., and Quinn, F. (2020). Interferon-gamma promotes iron export in human macrophages to limit intracellular bacterial replication. PLoS One 15, e0240949. 10.1371/journal.pone.0240949.

118. Temmerman, R., Vervaeren, H., Noseda, B., Boon, N., and Verstraete, W. (2006). Necrotrophic growth of Legionella pneumophila. Appl Environ Microbiol 72, 4323–4328. 10.1128/AEM.00070-06.

119. Archer, K.A., and Roy, C.R. (2006). MyD88-dependent responses involving toll-like receptor 2 are important for protection and clearance of Legionella pneumophila in a mouse model of Legionnaires’ disease. Infect Immun 74, 3325–3333. 10.1128/IAI.02049-05.

120. Chidiac, C., Che, D., Pires-Cronenberger, S., Jarraud, S., Campese, C., Bissery, A., Weinbreck, P., Brun-Buisson, C., Sollet, J.P., Ecochard, R., et al. (2012). Factors associated with hospital mortality in community-acquired legionellosis in France. Eur Respir J 39, 963–970. 10.1183/09031936.00076911.

121. Kostic, A., Cukovic, K., Stankovic, L., Raskovic, Z., Nestorovic, J., Savic, D., Simovic, A., Prodanovic, T., Zivojinovic, S., Andrejevic, S., et al. (2022). The Different Clinical Courses of Legionnaires’ Disease in Newborns from the Same Maternity Hospital. Medicina (Kaunas) 58. 10.3390/medicina58091150.

122. Levy, M.L., Le Jeune, I., Woodhead, M.A., Macfarlaned, J.T., Lim, W.S., and British Thoracic Society Community Acquired Pneumonia in Adults Guideline, G. (2010). Primary care summary of the British Thoracic Society Guidelines for the management of community acquired pneumonia in adults: 2009 update. Endorsed by the Royal College of General Practitioners and the Primary Care Respiratory Society UK. Prim Care Respir J 19, 21–27. 10.4104/pcrj.2010.00014.

123. Wee, B.A., Alves, J., Lindsay, D.S.J., Klatt, A.B., Sargison, F.A., Cameron, R.L., Pickering, A., Gorzynski, J., Corander, J., Marttinen, P., et al. (2021). Population analysis of Legionella pneumophila reveals a basis for resistance to complement-mediated killing. Nat Commun 12, 7165. 10.1038/s41467-021-27478-z.

124. Feeley, J.C., Gibson, R.J., Gorman, G.W., Langford, N.C., Rasheed, J.K., Mackel, D.C., and Baine, W.B. (1979). Charcoal-yeast extract agar: primary isolation medium for Legionella pneumophila. J Clin Microbiol 10, 437–441. 10.1128/jcm.10.4.437-441.1979.

125. Merriam, J.J., Mathur, R., Maxfield-Boumil, R., and Isberg, R.R. (1997). Analysis of the Legionella pneumophila fliI gene: intracellular growth of a defined mutant defective for flagellum biosynthesis. Infect Immun 65, 2497–2501. 10.1128/iai.65.6.2497-2501.1997.

126. Abshire, C.F., Dragoi, A.M., Roy, C.R., and Ivanov, S.S. (2016). MTOR-Driven Metabolic Reprogramming Regulates Legionella pneumophila Intracellular Niche Homeostasis. PLoS Pathog 12, e1006088. 10.1371/journal.ppat.1006088.

