## Supplementary figures and images for "A nutritional immunity blockade controls extracellular bacterial replication in *Legionella pneumophila* infections"

### Supplemental Fig 1

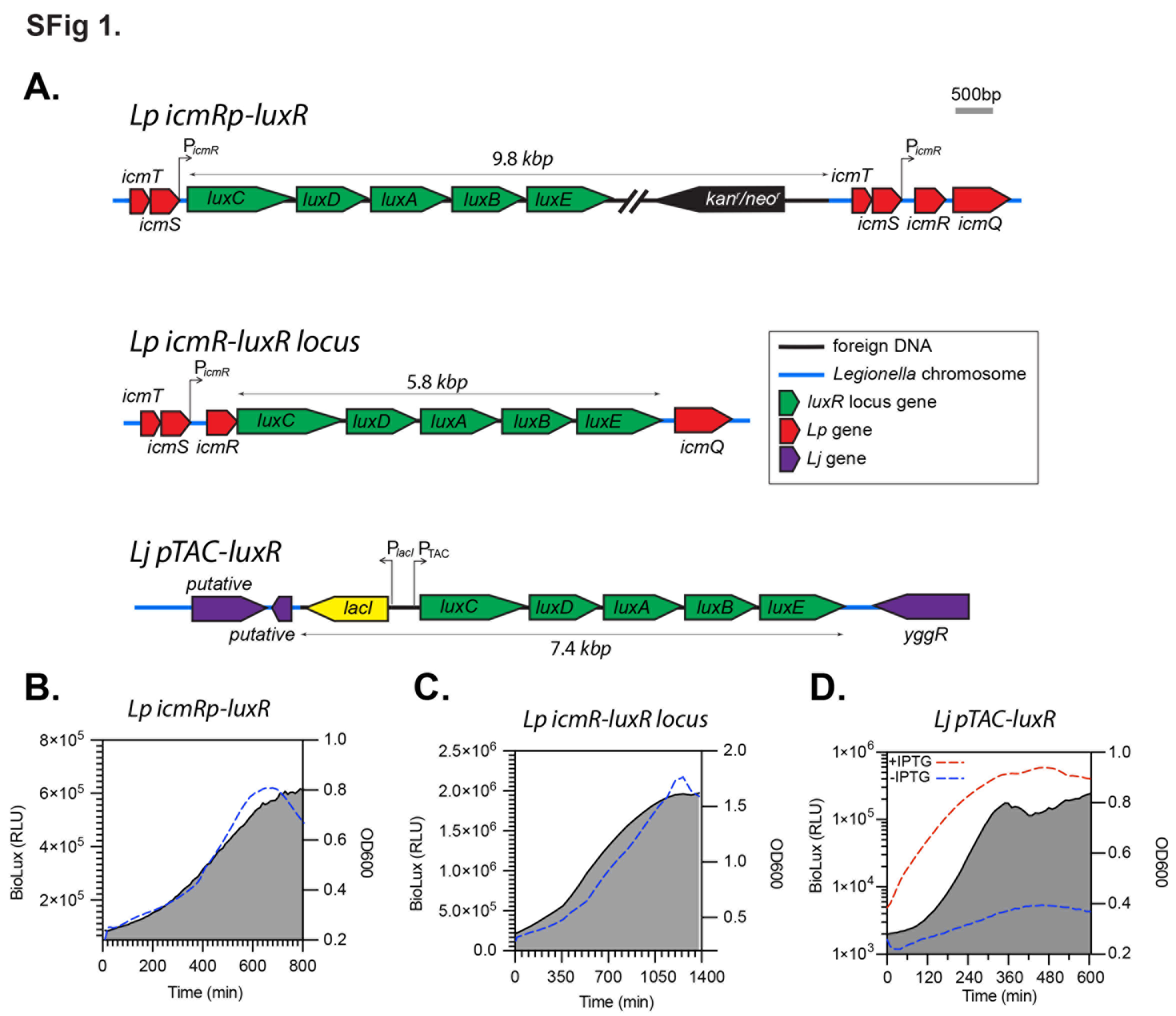

### Supplemental Fig 2

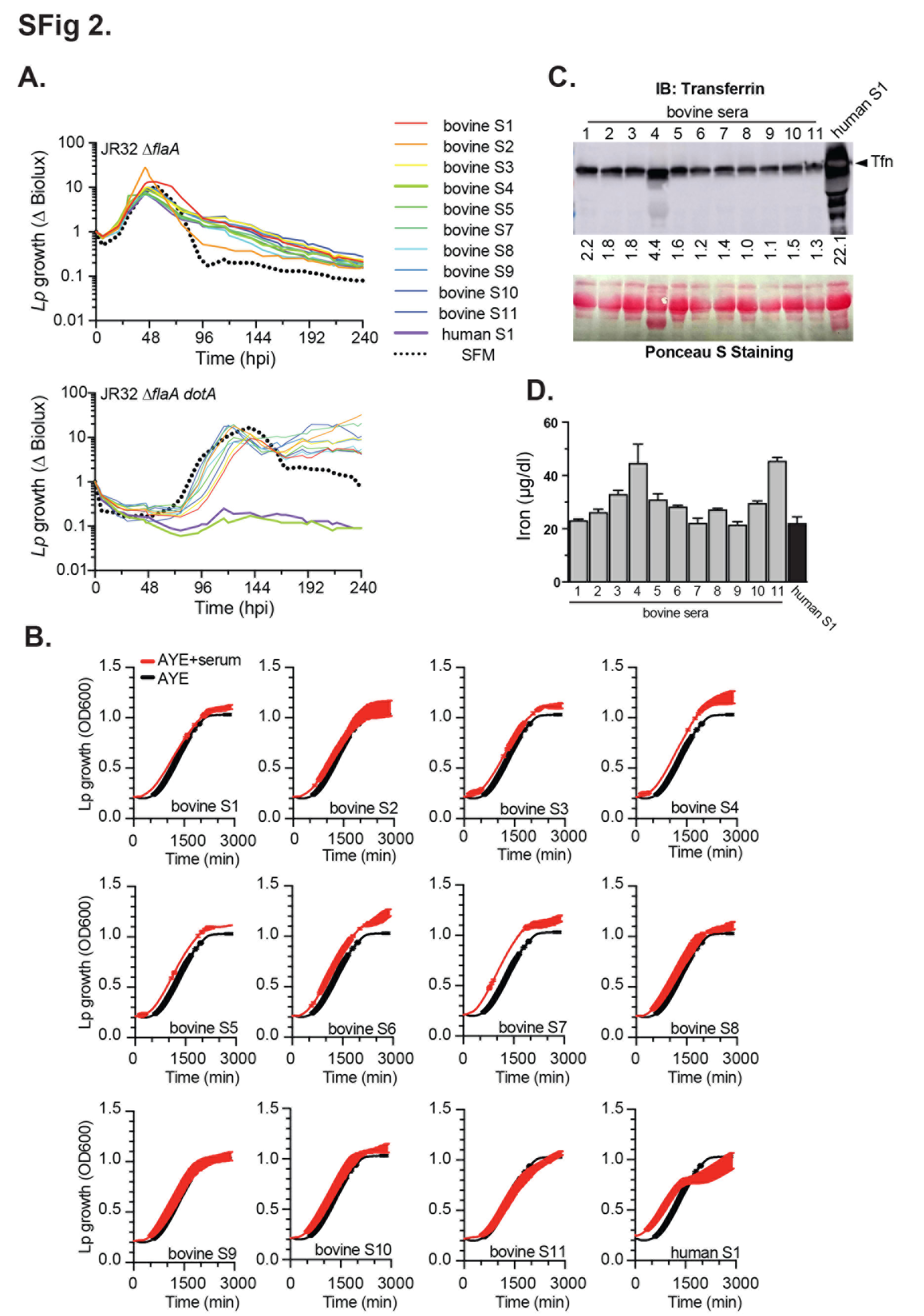

### Supplemental Fig 3

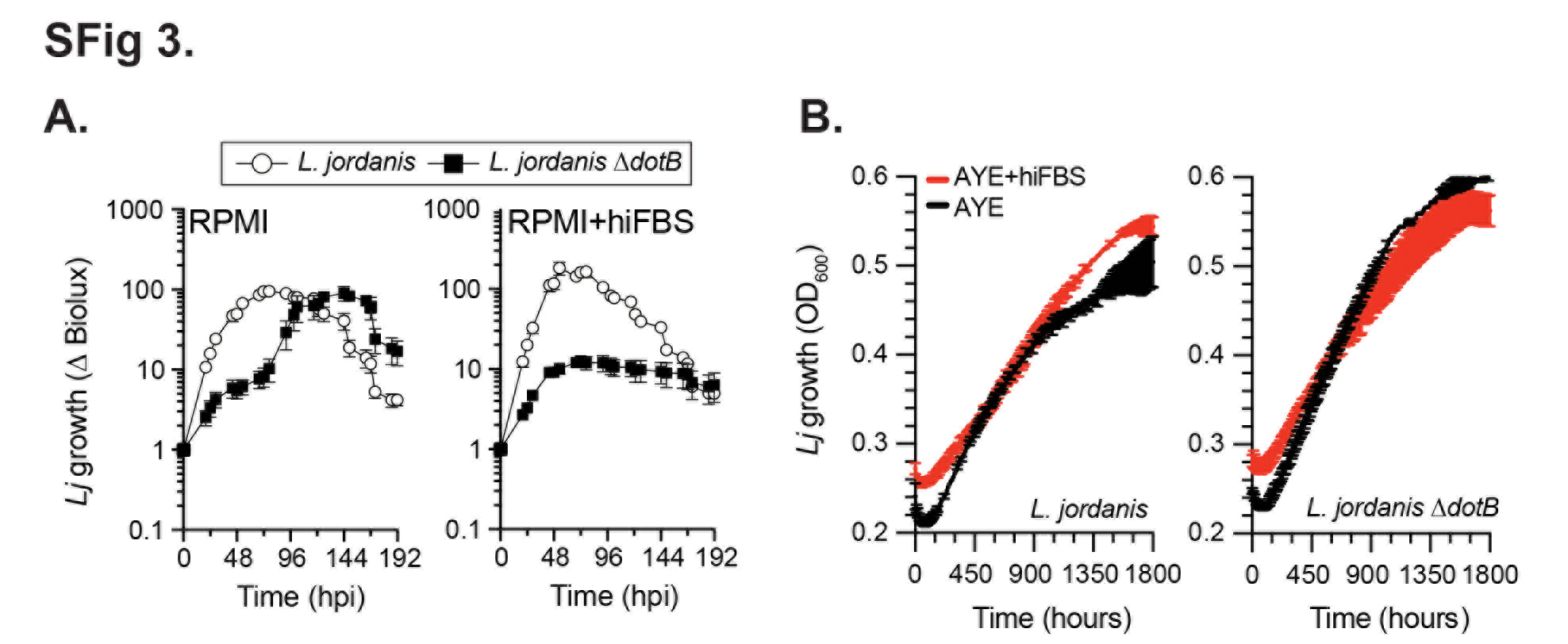

### Supplemental Fig 4

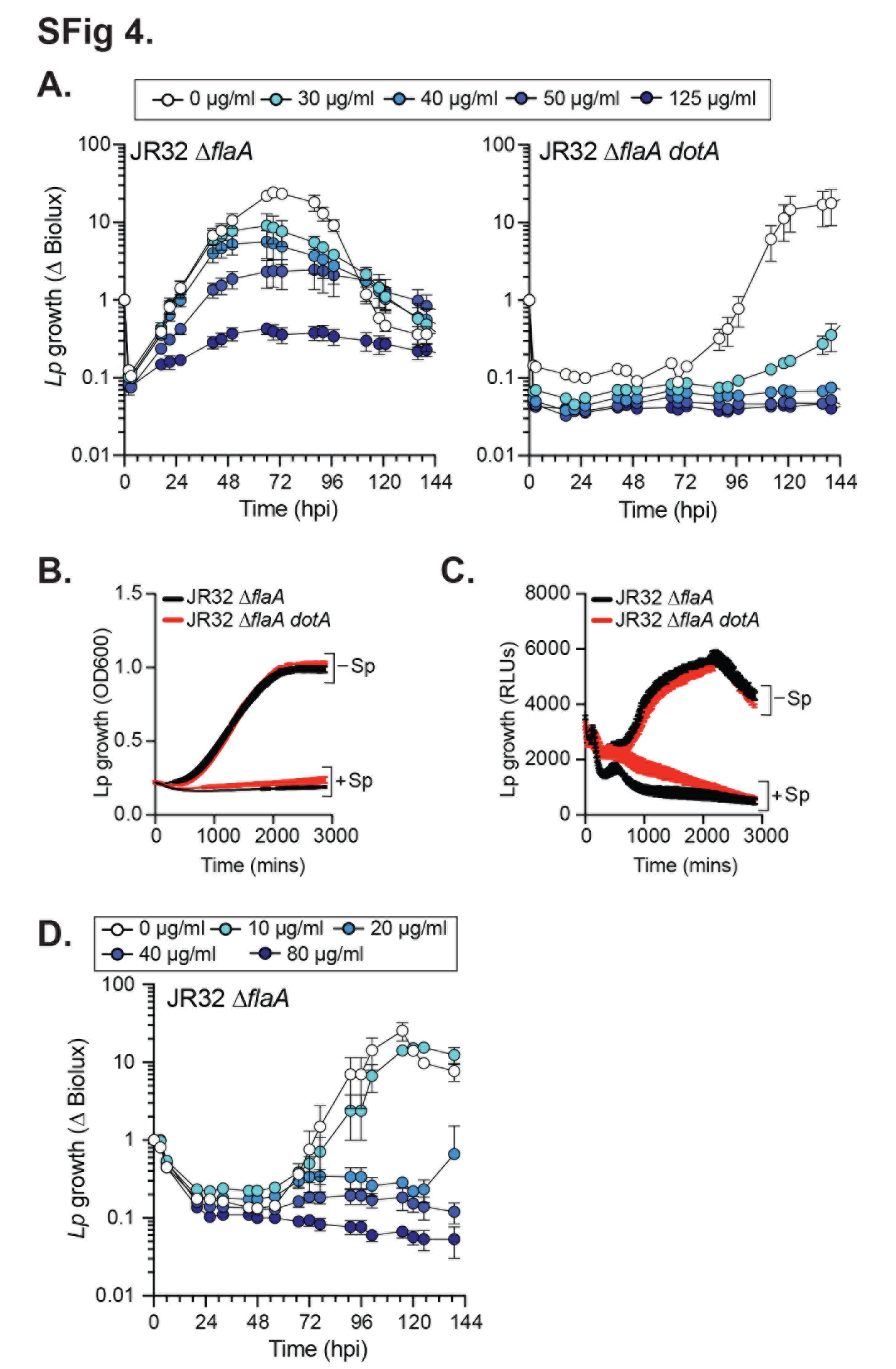

### Supplemental Fig 5

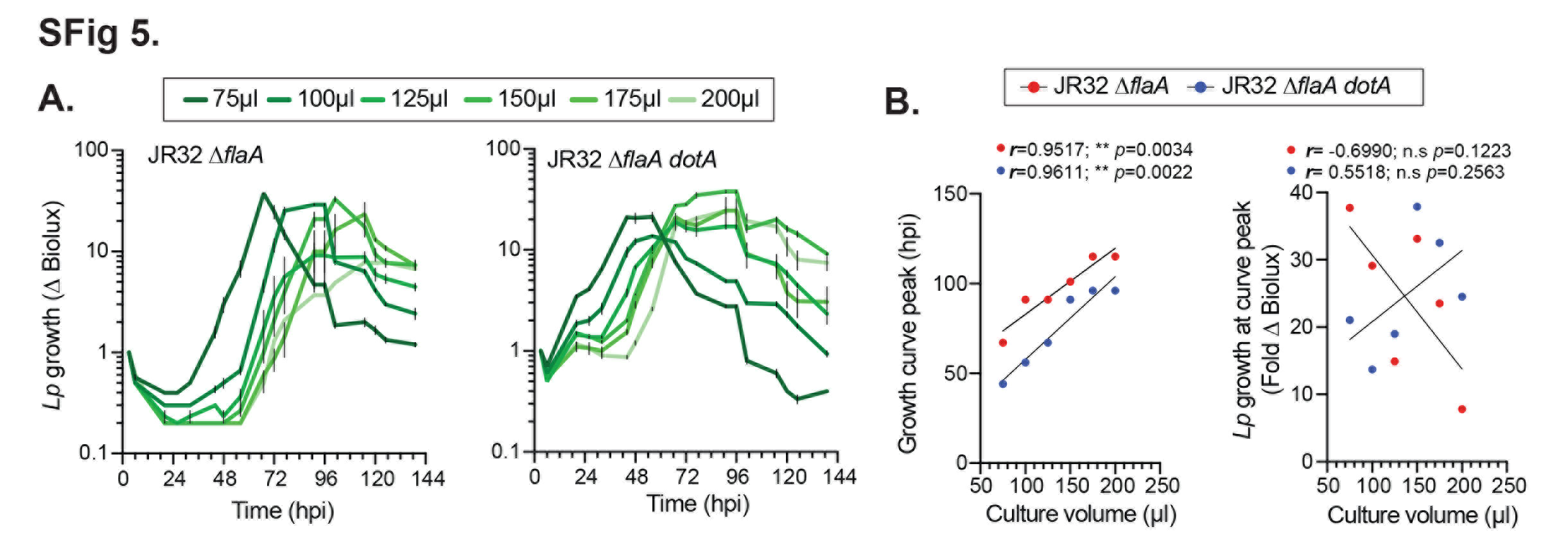

### Supplemental Fig 6

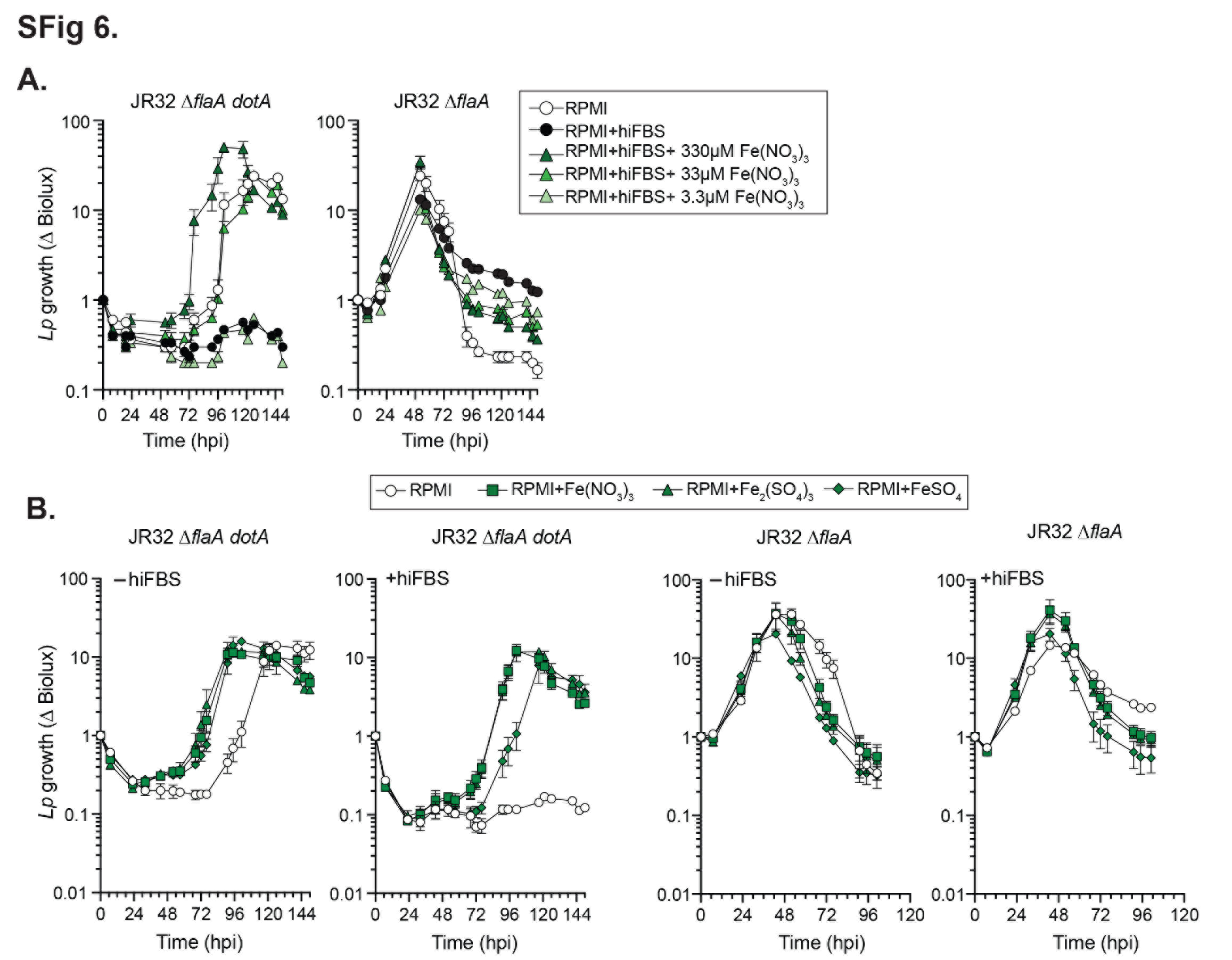

### Supplemental Fig 7

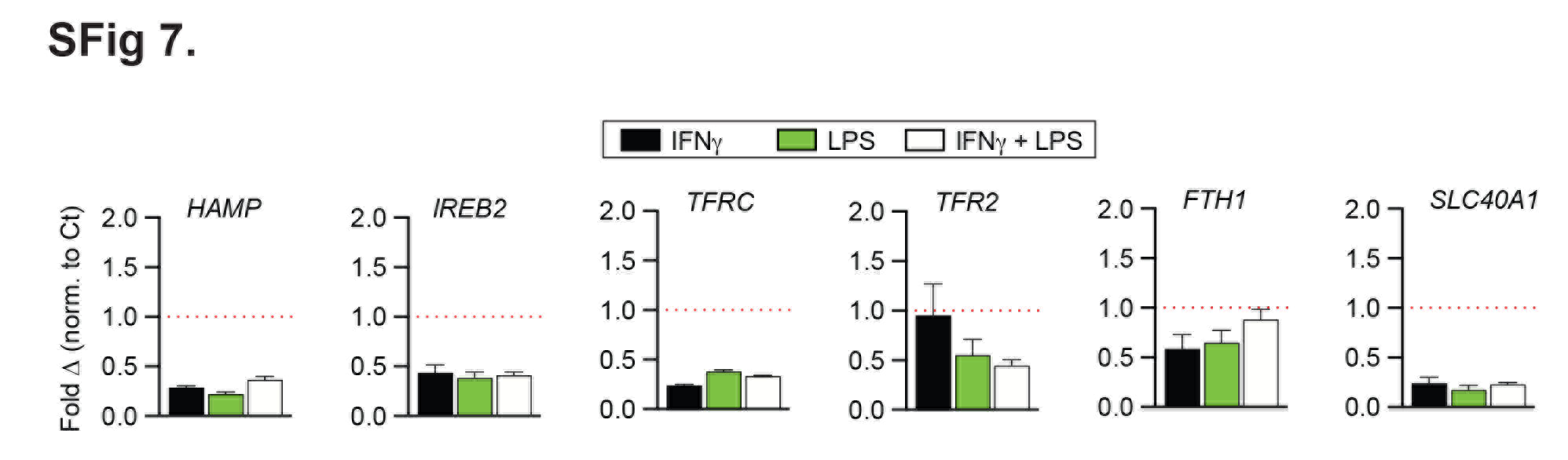
